# Microbiota supplementation with *Bifidobacterium* and *Lactobacillus* modifies the preterm infant gut microbiota and metabolome

**DOI:** 10.1101/698092

**Authors:** Cristina Alcon-Giner, Matthew J. Dalby, Shabhonam Caim, Jennifer Ketskemety, Alex Shaw, Kathleen Sim, Melissa Lawson, Raymond Kiu, Charlotte Leclaire, Lisa Chalklen, Magdalena Kujawska, Suparna Mitra, Fahmina Fardus-Reid, Gustav Belteki, Katherine McColl, Jonathan R. Swann, J. Simon Kroll, Paul Clarke, Lindsay J. Hall

**Affiliations:** Quadram Institute Bioscience, Norwich Research Park, Norwich, UK; Department of Medicine, Section of Pediatrics, Imperial College London, London, UK; Leeds Institute of Medical Research, University of Leeds, Leeds, UK; Department of Surgery and Cancer, Faculty of Medicine, Imperial College London, London, UK; Neonatal Intensive Care Unit, The Rosie Hospital, Cambridge University Hospitals NHS Foundation Trust, Cambridge, UK; Neonatal Intensive Care Unit, Norfolk and Norwich University Hospital, Norwich, UK; Norwich Medical School, University of East Anglia, Norwich, UK

## Abstract

Supplementation with members of the early-life microbiota or ‘probiotics’ is becoming increasingly popular to attempt to beneficially manipulate the preterm gut microbiota. We performed a large longitudinal study comprising two preterm groups; 101 orally supplemented with *Bifidobacterium* and *Lactobacillus* (Bif/Lacto) and 133 non-supplemented (Control) matched by age, sex, birth-mode, and diet. 16S rRNA metataxonomic profiling on stool samples (n = 592) indicated a predominance of *Bifidobacterium*, and a reduction of pathobionts in the Bif/Lacto group. Metabolic phenotyping found a parallel increase in fecal acetate and lactate in the Bif/Lacto group compared to the Control group, which positively correlated with *Bifidobacterium* abundance consistent with the ability of the supplemented *Bifidobacterium* strain to metabolize human milk oligosaccharides and reduced gut pH. This study demonstrates that microbiota supplementation can modify the preterm microbiome and the gastrointestinal environment to more closely resemble that of a full-term infant.

## Introduction

Infants born < 37 weeks gestation are defined as preterm, and account for 1 in 9 births globally (WHO, 2018). Compared to full-term infants preterm infants are more often born via Caesarean-section, have an underdeveloped immune system, receive numerous courses of antibiotics, and reside in neonatal intensive care units (NICUs); all of which disrupt the establishment of the early life gut microbiota (Gasparrini et al., 2019; Mulder et al., 2011; Walker, 2017). This altered gut microbial ecosystem has been linked to an increased risk of serious morbidity during the NICU stay, including necrotizing enterocolitis (NEC) (Shulhan et al., 2017) and late onset sepsis (LOS) (Pammi and Weisman, 2015), and later life health problems such as asthma and eczema (Been et al., 2014; Haataja et al., 2016).

Abnormal patterns of bacterial colonization are common in the preterm infant gut, which is dominated by genus containing potentially pathogenic bacteria (i.e. pathobionts) such as *Staphylococcus, Klebsiella, Escherichia*, and *Clostridium* (Dahl et al., 2018; Gasparrini et al., 2019). These infants are also characterized by a low abundance or absence of the beneficial *Bifidobacterium* and *Lactobacillus* that characteristic of the full-term infant gut (Dahl et al., 2018). Thus, interventions to ‘normalize’ the preterm gut microbiota are an attractive proposition to improve health and prevent disease in preterm infants.

Oral administration of commensal infant bacteria via probiotic (Hill et al., 2014) supplementation is one approach to encourage gut colonization by beneficial members of the early life microbiota. Systematic review and meta-analysis of randomised controlled trials and observational studies have reported that probiotic supplementation reduces NEC, sepsis, and all-cause mortality in preterm infants (AlFaleh and Anabrees, 2014; Dermyshi et al., 2017). However, one of the largest trials carried out in the UK found no evidence of benefit (Costeloe et al., 2016). Despite the outcome of previous systematic reviews and meta-analyses a 2018 survey of all 58 UK tertiary-level NICUs found only 10 (17%) were routinely using probiotics (Duffield and Clarke, 2019). While clinical studies have demonstrated the potential of probiotics to reduce NEC incidence, there are few that have also performed accompanying longitudinal microbiota profiling (often with relatively low numbers of infants(Esaiassen et al., 2018; Plummer et al., 2018; Watkins et al., 2019)) to determine the impact of supplementation on gut microbiota composition, and little or none that have examined the corresponding metabolome of preterm infants(Abdulkadir et al., 2016), nor included whole genome sequencing of probiotic bacteria isolated from the supplement used or from the recipient infants.

A recent clinical audit at the Norfolk and Norwich University Hospital NICU found rates of NEC fell from 7.5% to 3.1% and rates of late-onset sepsis fell from 22.6% to 11.5% when comparing the five years before and five years after the initiation of routine probiotic use with a combined *Bifidobacterium* and *Lactobacillus* supplement (P, 2019). Building on these important clinical observations, we aimed to explore the gut microbiota composition and faecal metabolome in these preterm infants receiving routine probiotic supplementation compared to preterm infants from NICUs not using probiotic supplementation.

Thus, we carried out an observational study comparing longitudinal samples from two cohorts of preterm infants; 101 orally supplemented with a combination of *Bifidobacterium* and *Lactobacillus* at the Norfolk and Norwich University Hospital NICU and 133 non-supplemented infants from NICUs not using probiotic supplementation. Cohorts were approximately matched by gestational age, sex, delivery method, sample collection time, and diet across the four tertiary-level NICUs. 16S rRNA gene profiling was used to determine the fecal bacterial composition (*n* = 592) and paired ^1^H nuclear magnetic resonance (NMR) spectroscopy was used to measure the metabolic content of the fecal samples, this included metabolites of microbial, host, and maternal origin (*n* = 157). To evaluate colonization potential whole genome sequencing was used to compare supplemented strains to isolates obtained from preterm infants, alongside *in vitro* studies to define underlying mechanisms governing preterm microbiota profiles.

## Results

### Study design

Fecal samples were collected from NICU-resident preterm infants receiving a daily oral supplementation containing *Bifidobacterium bifidum* and *Lactobacillus acidophilus* (Bif/Lacto group), and from a group of similarly aged preterm infants (Control group) from three other NICUs that did not offer routine supplementation. This observational study design avoided cross-contamination amongst the study groups, which has been reported previously in other probiotic studies where study groups reside within the same NICU (Costeloe et al., 2016; Deshpande et al., 2016; Henderickx et al., 2019; Hickey et al., 2014). Samples were collected corresponding to four time points at 0-9, 10-29, 30-49, and 50-99 days of age from birth (Fig. 1a).

**Figure 1:**
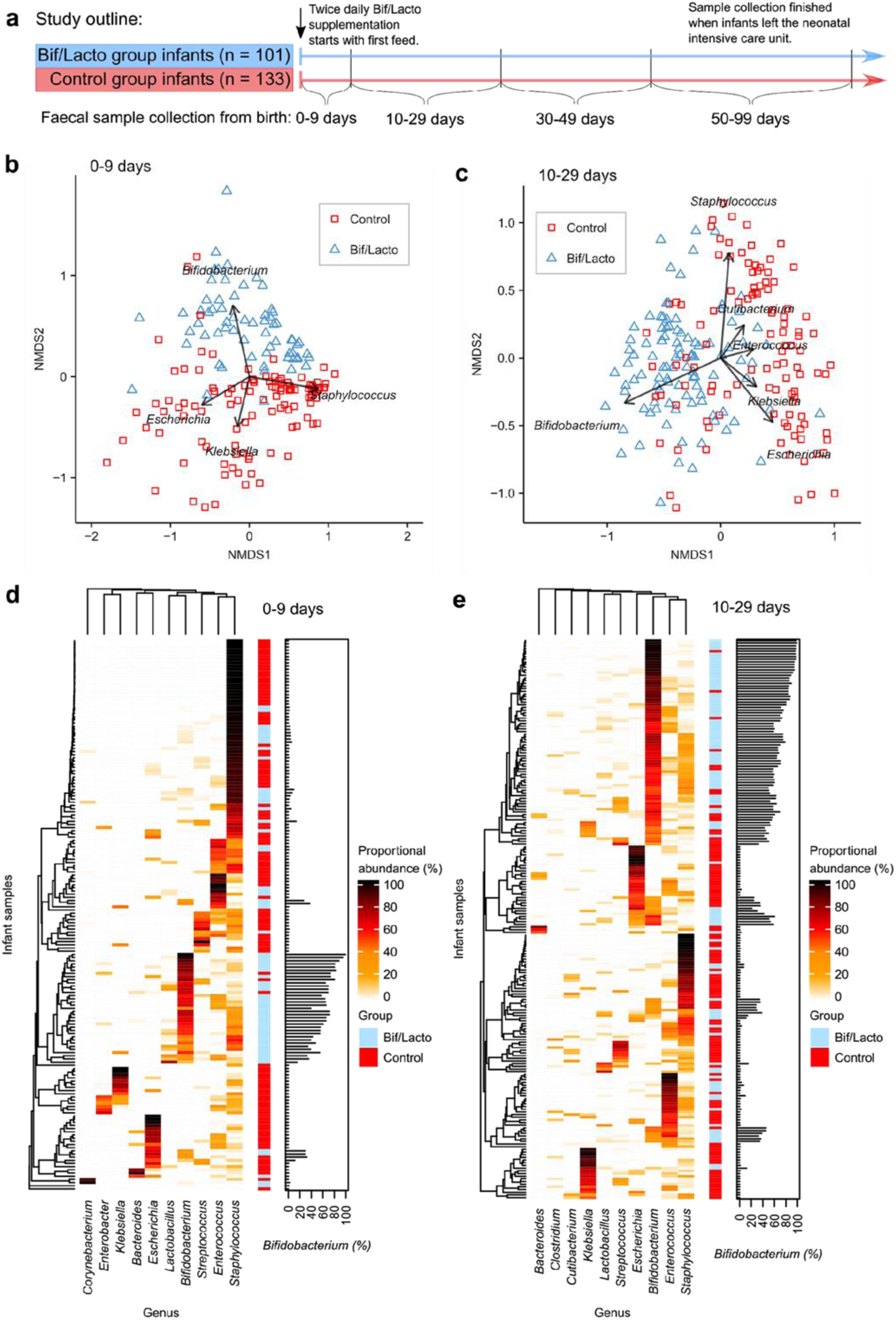
Premature infant gut microbiota clustering and genus composition. **a**, Study outline and sample collections times. Infloran supplementation was given until 34 weeks post-menstrual age, with the exception of very low birth weight infants (<1500 g) who received it until discharge. Control group was not given supplementation. **b**, Infant fecal microbiota similarity at 0-9 days and **c**, at 10-29 days shown using NMDS (Non-metric multidimensional scaling) analysis clustered with a Bray-Curtis dissimilarity. Arrows indicate bacterial genera driving the separation of points on the NMDS plots. **d**, Heatmaps showing the ten genera with highest proportional abundance at 0-9 days and **e**, at 10-29 days of age. Heatmap rows were clustered by total microbiota similarity using Bray-Curtis dissimilarity and the columns clustered by genera that occur more often together. Side bar plots show the proportional abundance of *Bifidobacterium* in each sample.

### Supplementation with early life microbiota members influences preterm gut microbiota composition

The preterm gut is typically dominated by pathobionts such as *Enterobacter, Escherichia*, and *Klebsiella*^*14*^. We sought to determine whether preterm infants supplemented with *Bifidobacterium and Lactobacillus*, bacterial species associated with a healthy term infant gut, showed a modified preterm microbiota profile. Fecal metataxonomic bacterial composition was determined by 16S rRNA gene sequencing. Genus level clustering of samples using non-metric multidimensional scaling (NMDS) indicated clear variation in the microbiota profiles between Bif/Lacto supplemented infants and Controls (Fig. 1b-c; Supp Fig. 1a-b). The microbiota composition of Bif/Lacto and Control sample differed significantly at each of the four time points (PERMANOVA, P < 0.01). The clustering of the Bif/Lacto group was driven by the genus *Bifidobacterium*, while the genera driving the clustering of the Control group included *Staphylococcus, Escherichia*, and *Klebsiella* (Fig. 1b-c; Supp Fig 1a-b). Notably, while hospital NICUs may differ in their ‘environmental’ microbiota in ways that may influence infant colonization, NMDS and PERMANOVA tests showed no differences in the microbiota composition between infant samples from the three Control hospital NICUs (Supp Fig. 2a-d).

We also examined the ten most abundant genera by relative abundance at each time point using Bray-Curtis dissimilarity (Fig. 1d-e; Supp Fig. 1c-d), showing infant samples clustered into six main groups based on a single dominant bacterial genus; *Bifidobacterium, Escherichia, Enterococcus, Klebsiella, Staphylococcus*, or *Streptococcus.* These data indicate that the introduction of *Bifidobacterium* promote changes in the composition of the preterm gut microbiota.

### Oral Bif/Lacto supplementation influences bacterial genus abundance and bacterial diversity

We sought to further define the genus composition based on relative abundance and diversity measures underlying these changes in microbiota composition. *Bifidobacterium* dominated the microbiota of the Bif/Lacto group with high relative abundance at all time points compared to the Control group (Fig. 2a-c). This indicated ‘efficient’ colonization by the supplemented strain, or ecosystem re-modeling to encourage colonization of other *Bifidobacterium* spp. Surprisingly, *Lactobacillus* was only detected in a minority of infants, but with a higher relative abundance in Bif/Lacto infants compared to the Control group at all time points (Fig. 2d), which may indicate a more transient and limited colonization potential for this strain. The relative abundance of bacteria such as *Klebsiella, Escherichia*, and *Enterobacter* was lower in Bif/Lacto infants compared to Control infants at earlier time points 0-29 days of age (Fig. 2e-g), with *Klebsiella* still lower at 30-99 days of age (Fig. 2e). *Staphylococcus* was initially abundant in both groups, but rapidly decreased as the infants aged (Fig. 2a-b; Supp Fig. 6f) (Tauchi et al., 2019). Species level analysis of the 16S rRNA data matched the *Staphylococcus* present to *S. epidermidis* and *S. haemolyticus*, bacterial residents on the skin, indicating that these originate from initial colonization of skin-associated bacteria (Supplementary Fig. 6g-h). Indeed, the skin-associated commensal *Cutibacterium* was also found in higher relative abundance in Control infants at 10-29 days of age (Fig. 2h). These data suggest that the oral supplementation impacts the microbial colonization patterns, displacing other potentially pathogenic bacteria more typical of the preterm gut.

**Figure 2:**
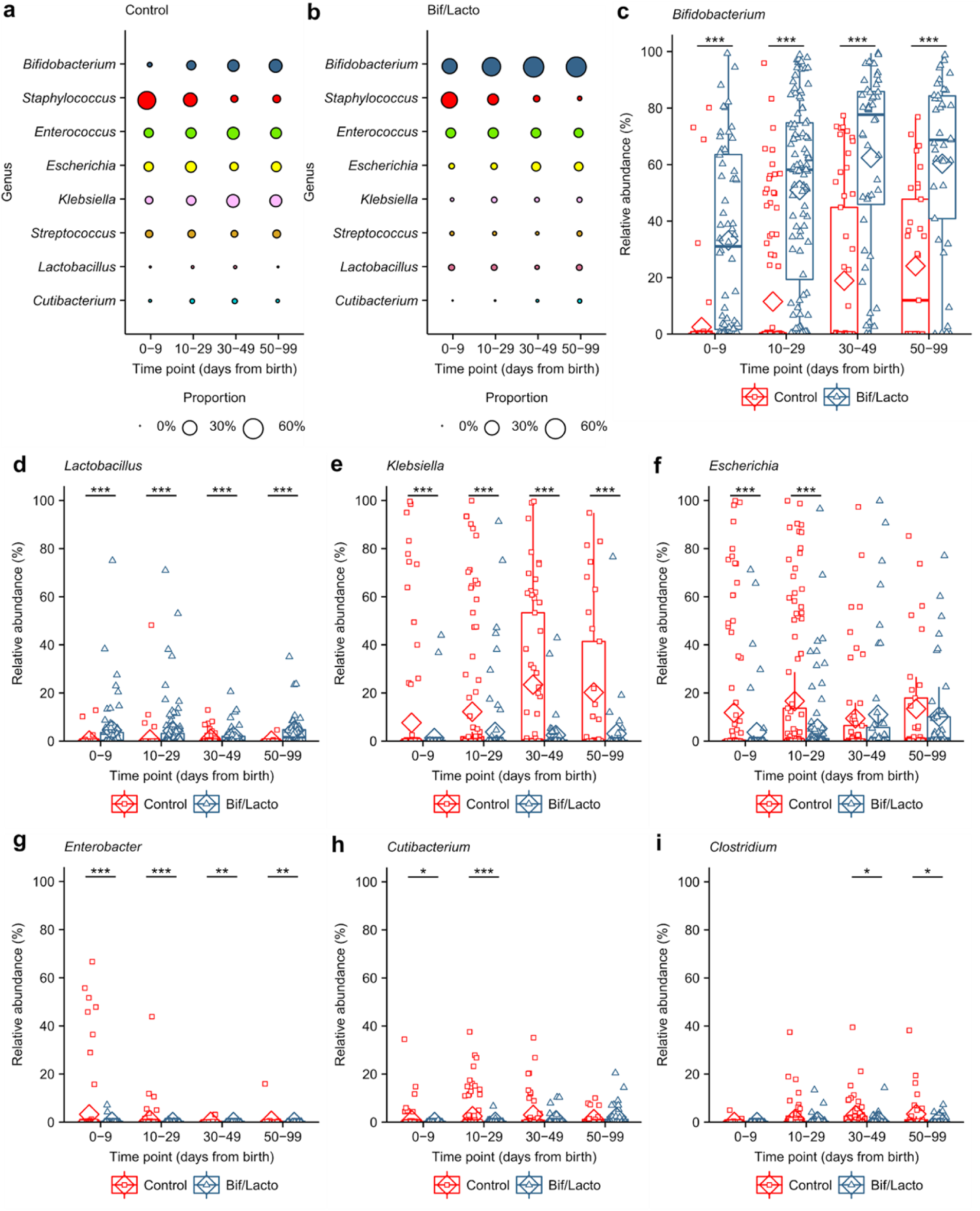
Genus abundance between Bif/Lacto and Control groups. **a-b**, Bubble plots show the mean group abundance of the common bacterial genera at each time point in the Control group and the Bif/Lacto group. **c**, *Bifidobacterium* abundance at each time point. Relative abundance of **d**, *Lactobacillus*, **e**, *Klebsiella*, **f**, *Escherichia*, **g**, *Enterobacter*, **h**, *Cutibacterium*, and **i**, *Clostridium*. Individual points highlight individual infant samples, diamonds indicate the group mean, box plots show group median and interquartile range. Asterisks represent *p* values: ***P < 0.001.

When examining diversity measures (Shannon and Inverse Simpson diversity), values were initially higher at 0-9 days in Bif/Lacto compared to Control infants (Supp Fig. 3b-c), although the abundance of *Bifidobacterium* was not correlated with the number of bacterial genera detected (Supp Fig. 3d). At later time-points (i.e. 30-99 days of age, Fig Supp Fig. 3b-c), the diversity values of the Bif/Lacto were lower than the Control group, which may correlate with the increasing *Bifidobacterium* abundance above 50% (Supp Fig. 3e-f, h-i). These data suggest there is a diversity ‘tipping-point’ in response to dominance of *Bifidobacterium* as the major early life member.

### External factors including gestational age, birth weight, and antibiotics negatively affect *Bifidobacterium* abundance in Bif/Lacto infants

Previous studies have indicated that factors, such as gestational age and antibiotics (Korpela et al., 2018), significantly influence the developing early life gut microbiota; with preterm infants representing an infant cohort overexposed to potential microbiome-modulating factors. Focusing on *Bifidobacterium* as the dominant bacteria in the Bif/Lacto group, we noted that infants with a birth weight ≥1000 grams showed higher relative abundance of *Bifidobacterium* at 0-29 days (Fig. 3a). This was also the case at 10-29 days of age in infants born at a gestational age ≥28 weeks. (Fig. 3b). Furthermore, birth weight and gestational age were closely correlated (Fig. 3c) and correlated inversely with length of NICU stay (Supp Fig. 4e-f), indicating that the underdeveloped preterm gut may not represent an optimal niche for *Bifidobacterium* colonization. Higher *Bifidobacterium* proportions in Control infants with birth weights ≥1000 g compared to those of <1000 g supports this hypothesis (Supp Fig. 4a). There was no difference in length of stay in NICU between Bif/Lacto and Control infants (Supp Table 1, Supp Fig. 5e).

**Figure 3:**
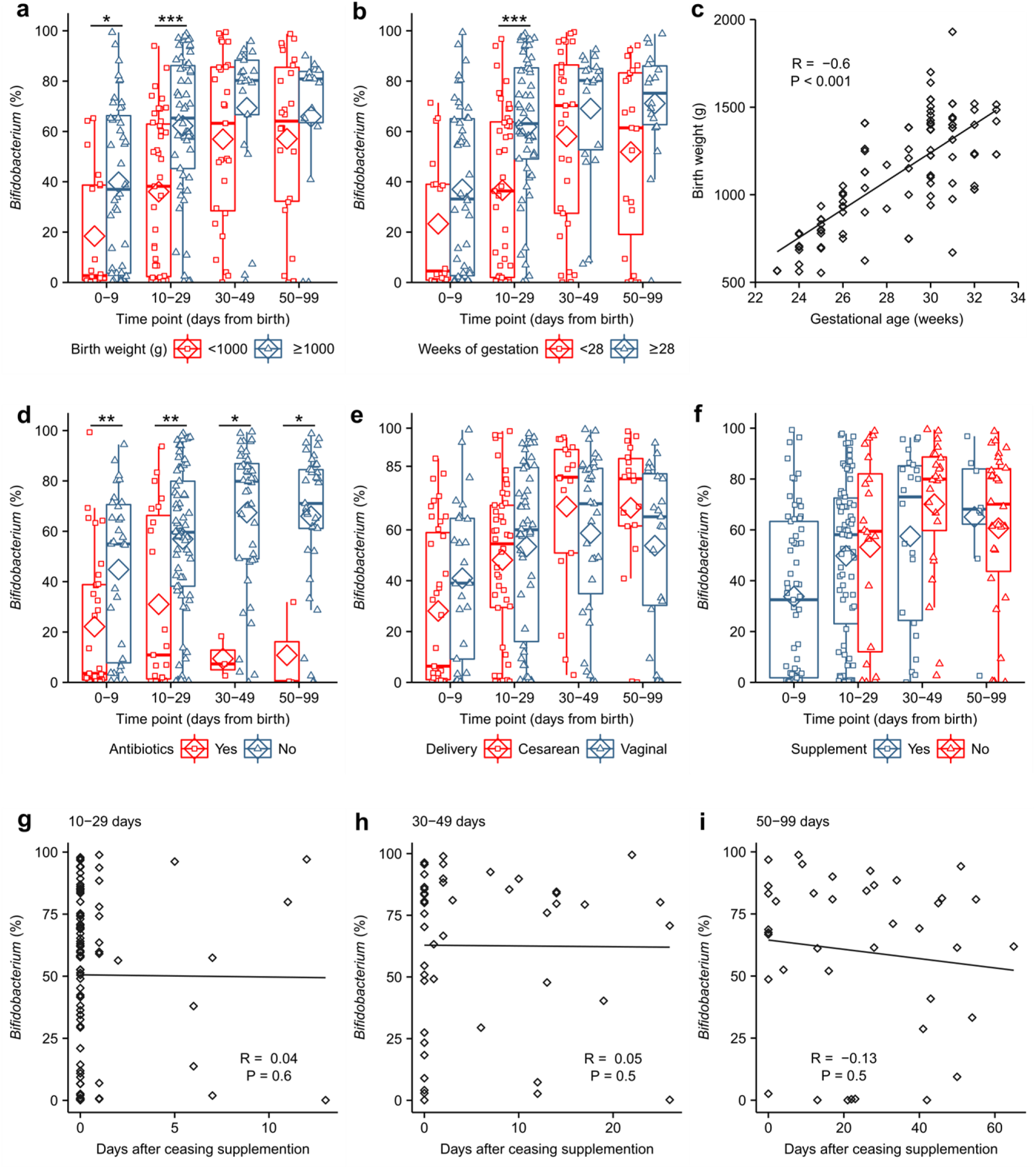
Effects of birthweight, antibiotic use, delivery mode and *Bifidobacterium* colonisation on Bif/Lacto infants. **a**, *Bifidobacterium* abundance between very-low birth weight (<1000 g) and low birth weight (>1000 g) **b**, *Bifidobacterium* abundance between infants with very low gestational age (<28 weeks) and low gestational age (≥28 weeks). **c**, Birth weight in grams correlated with gestational age in weeks. **d**, *Bifidobacterium* abundance in infants receiving antibiotics at the time of sample collection. **e**, *Bifidobacterium* abundance in infants delivered by caesarean and vaginal birth. **f**, *Bifidobacterium* abundance in infants still receiving or no-longer receiving supplementation. **g**-**i** Correlation between *Bifidobacterium* abundance and days after ceasing receiving supplementation. Asterisks represent *p* values: *P < 0.05, **P < 0.01, ***P < 0.001.

Preterm infants receive numerous antibiotics over the course of their NICU stay. Within the Bif/Lacto infants *Bifidobacterium* abundance was lower in infants currently being treated with antibiotics at all time points compared to those not receiving antibiotics, indicating antibiotic susceptibility of this genus (Fig. 3d). In contrast, the relative abundance of *Staphylococcus, Klebsiella*, and *Escherichia* was unchanged in infants receiving antibiotics, suggesting these were resistant to antibiotic treatment (Supp Fig. 4i-k).

Emergency Caesarean sections for maternal or fetal indications account for a large number of preterm births, and previous studies have indicated that Caesarean-section delivery can directly interrupt the transfer of maternal microbes (e.g. *Bifidobacterium*) to infants (Dominguez-Bello et al., 2010; Shao et al., 2019). Surprisingly, there was no significant difference in the relative abundance of *Bifidobacterium* within the Bif/Lacto group in infants born by vaginal or cesarean birth (Fig. 3e). Gestational age, current antibiotic treatment, and delivery method did not significantly alter *Bifidobacterium* proportions in Control infants, however the low abundance of this bacteria in this cohort make robust statistical analysis difficult (Supp Fig. 4b-d).

In Bif/Lacto infants, supplementation ceased when infants reached a post-conceptual age of 34 weeks. However, no reduction was observed in the relative abundance of *Bifidobacterium* in samples collected from these infants after oral supplementation had ceased (Fig. 3f), with proportions maintained for up to 60 days (Fig. 3g-i). This indicates potentially longer-term colonization of the supplemented strain in these infants.

Diet is proposed to be one of the major factors modulating the early life microbiota, with significantly differences between formula and breast-fed infants (Timmerman et al., 2017). Unusually, almost all infants recruited to this study were fed either their own mothers’ breast milk, or their mothers’ breast milk and donor breast milk in combination, or breast milk supplemented with preterm cows’ milk-based formula. Only a very small number of infants were exclusively formula fed (i.e. 4 out of 234), which may explain our findings that the relative abundance of *Bifidobacterium* was not statistically significantly affected by diet within both the Bif/Lacto or Control infant cohorts (Supp Fig. 4g-h). There were no differences in the microbiota composition in either Bif/Lacto infants or Control infants between those fed mothers breast milk compared to those fed a combination of mothers’ breast milk and donor breast milk (Supp Fig. 5a-d).

### Bif/Lacto infants shown distinct changes in species composition over time including prolonged colonization of the *Bifidobacterium bifidum* Infloran strain, which correlates with human breast milk metabolism and routine supplementation

Our data so far indicated a dominance of *Bifidobacterium* in the Bif/Lacto cohort, and poor colonization ability of *Lactobacillus* in the preterm gut. To probe this with greater resolution, we compared the abundance of species present within these two genera. *B. bifidum* was highly abundant in Bif/Lacto infants, while only being abundant in 2/133 Control infants (Fig. 4a). *Bifidobacterium breve* was also more abundant in the Bif/Lacto supplemented group (Supp Fig. 6a), with *Bifidobacterium longum* present in a small number of infants from both groups (Supp Fig. 6b). *B. bifidum* relative abundance declined with increasing infant age (Fig. 4a), with concurrent increases in *B. breve* (Supp Fig. 6a). As *B. breve* coexisted with, rather than replaced (Supp Fig. 6i) *B. bifidum* this suggests close species interactions; potentially via metabolite cross-feeding. In contrast, while the *Lactobacillus acidophilus* Infloran strain (for genome analysis see Supp Fig. 7d-e) was more prevalent in Bif/Lacto infants (Supp Fig. 6d), abundance decreased to zero within days after cessation of supplementation (Supp Fig. 6e-h), indicating inefficient colonization of this species.

**Figure 4:**
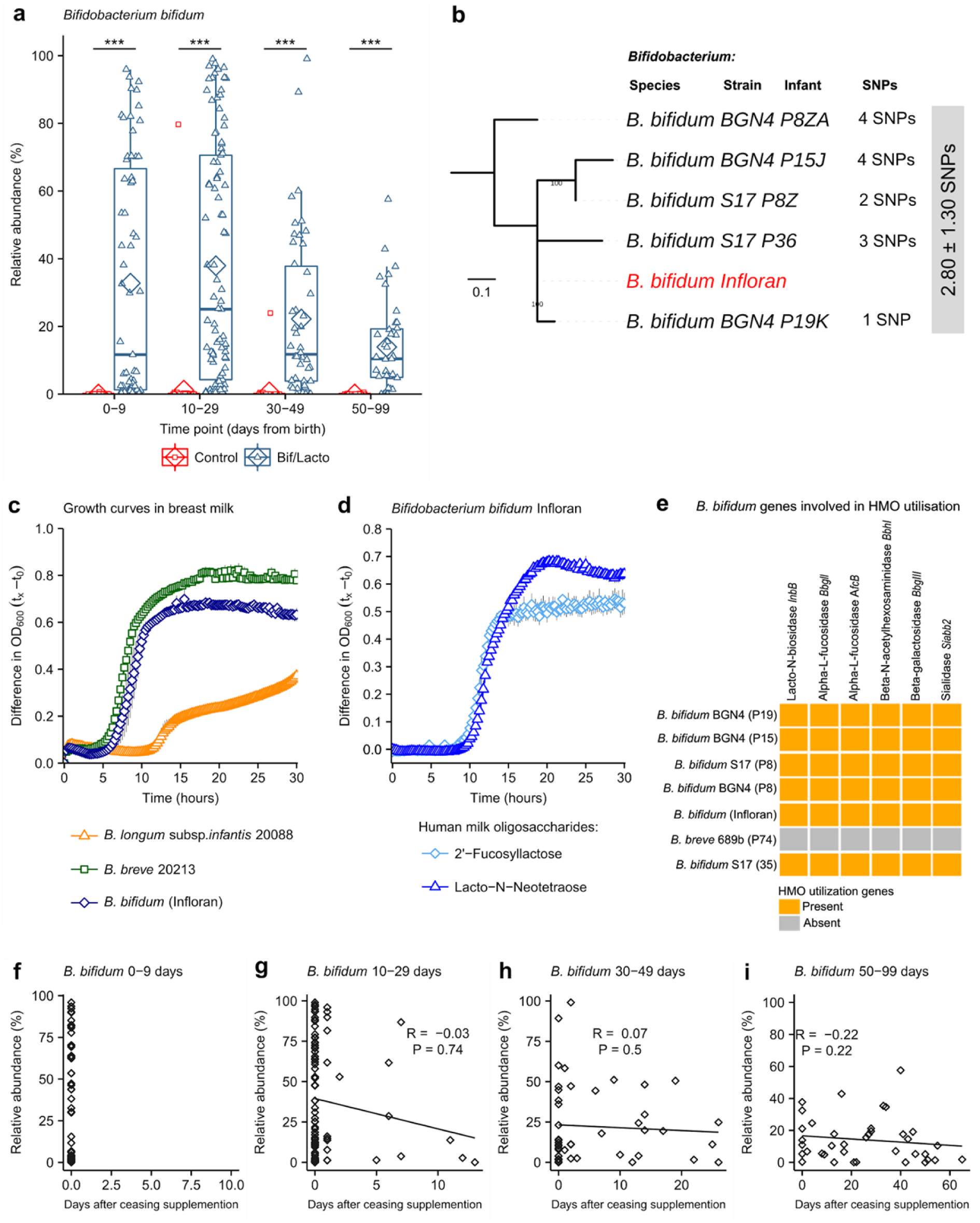
Comparison of *B. bifidum* genomes, and phenotypic characterisation of *B.bifidum* Infloran strain. **a**, *Bifidobacterium bifidum* abundance in Bif/Lacto and Control group infants. **b**, Mid-point rooted maximum-likelihood tree based on 12 SNPs called via reference-based approach (strain Infloran as the reference genome) from 5 *Bifidobacterium bifidum* genomes. Grey box denotes pairwise SNP distance between these 6 genomes. Data: mean ± S.D. **c**, Growth curves of *B. bifidum* Infloran, *B. breve* 20213, and *B. longum* subsp. *infantis* 20088, in whole human milk. **d**, Growth curves *B. bifidum* infloran in human milk oligosaccharides (HMO) Lacto-N-tetraose and 2-fucosyllactose **e**, Heat map representing *B. bifidum* genes involved in utilisation of human milk oligosaccharides. **f-i**, Correlation between *B.bifidum* abundance and days after ceasing receiving supplementation. Asterisks represent *p* values: ***P < 0.001.

Previous research studies have shown that colonization of the gut by probiotic bacteria may vary depending on the strains used, mode of administration, dose, and inclusion of prebiotics (Suez et al., 2019). To confirm putative bifidobacterial supplemented strain colonization (as described above), we performed whole genome sequencing-based analysis to compare the *B. bifidum* Infloran strain to nine *Bifidobacterium* isolates cultured from fecal samples from seven Bif/Lacto supplemented infants. Core-genome single nucleotide polymorphisms (SNPs) analysis indicated the five *B. bifidum* isolates were identical at 0 SNP difference (based on 87 core genes; Supp Fig. 7a). Reference-based genome-mapping of whole genome sequences of five *B. bifidum* genomes to *B. bifidum* Infloran strain (as reference genome, Fig 4b) indicated a near-identical similarity (mean pairwise SNPs: 2.80 ± 1.30), strongly suggesting they belong to the same bacterial strain (i.e. Infloran). Average nucleotide identity (ANI) analysis also supported these findings (100.00% nucleotide identity, Supp Fig 7b). These data support the elevated *B. bifidum* relative abundances in our 16S rRNA metataxonomic data (Fig. 4a), including after supplementation had finished in Bif/Lacto infants, indicating longer-term colonization of this strain (Fig. 4f-i).

*Bifidobacterium* represents a dominant genus in the full-term healthy breast-fed infant selectively fed by complex oligosaccharides (i.e. human milk oligosaccharides (HMOs)) within breast milk. However, the ability of *Bifidobacterium* to digest HMOs varies between species and strains of this genus (Gotoh et al., 2018). Thus, we analyzed *B. bifidum* genomes (our 5 isolates and Infloran strain) for the presence of genes involved in HMO utilization; all *B. bifidum* isolates contained specific genes involved in HMO utilization (Fig. 4e), and mucin degradation genes which may aid gut colonization (Supp Fig. 6c). Notably, growth curves in whole breast milk (Fig. 4c) confirmed that the *B. bifidum* Infloran strain utilized whole breast milk. Further phenotypic analysis indicated this strain was able to metabolize specific HMOs; 2-fucosyllactose (2’-FL) and Lacto-N-Neotetraose (LnNT), corresponding to genes *AfcA* and *BBgIII* respectively, encoding for extracellular enzymes involved in their utilization (Fig. 4d). Therefore, the ability to digest breast milk and HMOs in these predominantly breast milk fed infants, is likely to be a key factor driving high rates of *Bifidobacterium* colonization.

Bacterial strains used as probiotics commonly lack antibiotic resistance genes. However, the high levels of antibiotic usage in the NICU may reduce the colonization of supplemented strains in the preterm gut. Analysis of the *B. bifidum* Infloran strain genome indicated the presence of only the intrinsic *ileS* gene (associated with mupirocin resistance, Supp Table 2a). Minimum antibiotic concentration testing confirmed sensitivity to commonly prescribed antibiotics in NICUs (Supp Table 2b). These data are in agreement with the reduced relative abundance of *Bifidobacterium* in Bif/Lacto infants receiving antibiotics (Fig. 3d) However, by giving the supplement twice daily (up to 34 weeks post-conceptual age) this may have aided rapid re-establishment after antibiotic treatment and promoted relatively stable colonization.

### Infants receiving oral supplementation show differences in metabolomic profiles and lower fecal pH

Microbial metabolites are key molecules involved in microbe-microbe and microbe-host interactions (Peisl et al., 2018). To define the ‘functional’ impact of Bif/Lacto supplementation, ^1^H nuclear magnetic resonance (NMR) spectroscopy was used to characterize the metabolomes of a subset of paired fecal samples (75 from Bif/Lacto group, and 81 from Control group; all timepoints; N = 157). A principal component analysis (PCA) model (R^2^ = 53.6%) was built using these metabolic phenotypes and clear biochemical variation was observed between the Bif/Lacto and Control samples (Fig. 5a). Pairwise orthogonal projection to latent structures-discriminant analysis (OPLS-DA) models constructed for each time point confirmed these metabolic differences throughout the study period (p values < 0.01, Supp Fig. 7a). A covariate-adjusted-PLS-DA (CA-PLS-DA) model comparing the fecal profiles at all sampling points and adjusted for sampling age showed that infants in the Bif/Lacto group excreted greater amounts of the short chain fatty acid (SCFA) acetate (Fig. 5c) and lower amounts of the sugars 2’-FL, 3-fucosyllactose (3’-FL), arabinose, and trehalose compared to those in the Control group (Fig. 5e-h). Compared directly, fecal lactate was also higher in Bif/Lacto infants compared to Control infants (Fig. 5d). Notably, these differences observed were maintained throughout the study period.

**Figure 5:**
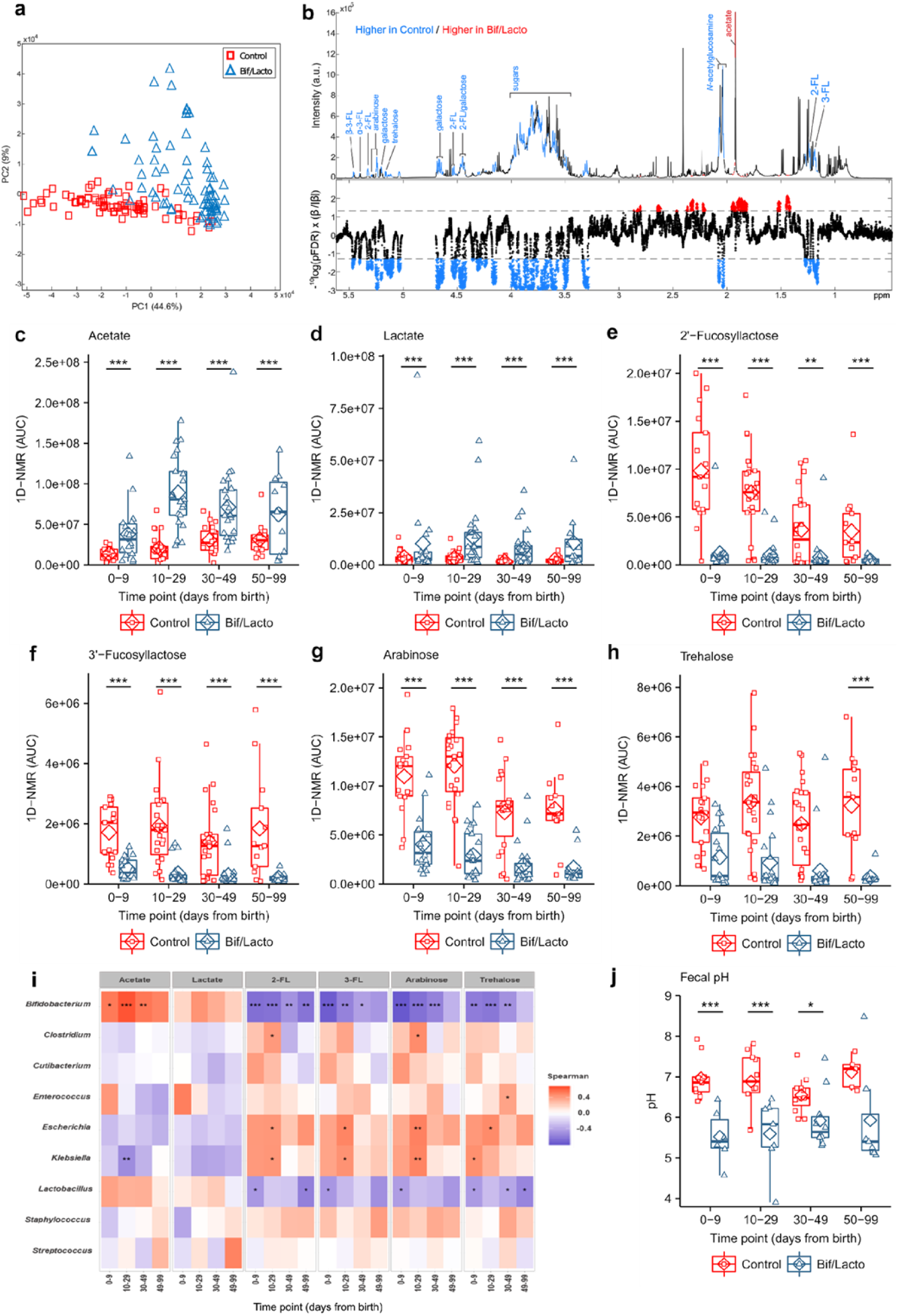
Metabolomic profiling of feces from the Bif/Lacto and Control Group using 1H NMR spectroscopy. **a**, Principal Component Analysis (PCA) scores plot comparing the fecal metabolic profiles of the Bif/Lacto and Control groups at all time points. **b**, Discriminatory metabolites that contribute to the covariate-adjusted projection to latent structures-discriminant analysis (CA-PLS-DA) model comparing the fecal metabolic profiles of the Bif/Lacto and Control infants adjusted for sampling age. Upper panel. Average 1H NMR spectrum from all samples indicating metabolites that are excreted in greater amounts by the Bif/Lacto infants (red) and those excreted in greater amounts by the control infants (blue). Lower panel. Manhattan plot showing P-values calculated for each variable in the multivariate model, corrected for multiple testing using the False Discovery Rate (allowing 5% false discoveries). Horizontal lines indicate cut-off values for the false discovery rate on the log_10_ scale. Blue points indicate metabolites significantly higher in the control feces and red points indicate those metabolites significantly higher in the Bif/Lacto feces. **c**, Relative acetate concentration. **d**, Relative lactate concentration. **e**, Relative 2’-fucosyllactose (2-FL) concentration. **f**, Relative 3’-fucosyllactose (3-FL) concentration. **g**, Relative arabinose concentration. **h**, Relative trehalose concentration. **i**, Spearman correlation heat map displaying main faecal metabolites (rows) versus the most abundant bacterial groups (columns). Red denotes positive correlation and blue denotes for negative correlation. **j**, Group faecal sample pH. Asterisks represent *p* values: \**p* < 0.05, \*\**p* < 0.01, \*\*\**p* < 0.001. Details of metabolites data for each sample investigated and pH measurements can be found in Supplementary Tables 8 and 9.

The relative abundance of *Bifidobacterium* was found to be significantly positively associated with fecal acetate and negatively associated with fecal 2’-FL, 3’-FL, arabinose, and trehalose (Fig 5i). Acetate and lactate are known metabolic by-products of *Bifidobacterium*^*21*^, while 2’-FL and, 3’-FL are common components of HMOs, with certain *Bifidobacterium* strains (including Infloran Fig 4e) able to selectively metabolize these breast milk components^22^. These results indicate that the higher relative abundance of *Bifidobacterium* in Bif/Lacto infants correlates with greater HMO metabolism, and with acetate and lactate generated as major end products.

To determine the impact of increased acetate and lactate on the infant gut environment, fecal pH was measured in a subset of infants (n = 74). At 0-9 days of age fecal pH was 5.5 (SD=0.7) in Bif/Lacto infants compared to pH 7.0 (SD=0.5) in Control infants. These differences in fecal pH remained throughout the study (Fig. 5j) and fecal pH was significantly negatively correlated with fecal acetate and lactate (Supp Fig. 8c-d) and the relative abundance of *Bifidobacterium* (Supp Fig. 8e). Comparing the relative bacterial abundance at species level, *B. bifidum* had a stronger negative correlation with fecal pH and positive correlation with fecal acetate and lactate (Supp Fig 9a-c) compared to *B. breve*, the other main *Bifidobacterium* species present (Supp Fig. 9d-f). Metabolomic analysis on bacterial culture supernatant confirmed the strong acetate producing ability of the supplemented strain *B. bifidum* (Supp Fig. 9g).

## Discussion

Our results show that preterm infants supplemented with *B. bifidum* and *L. acidophilus* contain a fecal microbiota composition and environment more similar to a healthy full-term breast-fed infant. Crucially, we determined that certain clinical practices and relevant external factors may positively or negatively influence the ability of introduced strains, particularly *Bifidobacterium*, to successfully colonize and establish a niche within the preterm infant gut microbiota.

As diet is a major driver of microbiota diversity, particularly the strong relationship between breastmilk and *Bifidobacterium* abundance, we were careful to match our study cohorts (Kirmiz et al., 2018). Although both groups of preterm infants received high rates of breast milk, via maternal or donor milk, the low abundance of *Bifidobacterium* found in Control infants indicates breast milk consumption itself (without supplementation), was not sufficient to encourage high rates of ‘natural’ *Bifidobacterium* colonization. Maternal to infant transmission of *Bifidobacterium* occurs in term infants (Ayechu-Muruzabal et al., 2018; Milani et al., 2015; Shao et al., 2019), however the NICU environment and antibiotic treatment may limit establishment of parental *Bifidobacterium*, leaving infants susceptible to colonization by hospital-environmental bacteria (Gasparrini et al., 2019; Shao et al., 2019; Zou et al., 2018). For the Bif/Lacto group, the combination of supplementation of early life microbiota members and a known prebiotic food source i.e. breast milk and HMOs, likely aided colonization (Kirmiz et al., 2018). Crucially, this symbiotic approach was successful because the ‘right’ bacterial strain was matched to the appropriate nutritional environment. In this case a *B. bifidum* strain with the genetic potential to metabolize HMOs, and phenotypically shown to use these early life dietary sources for growth. Interestingly, previous studies show *B. bifidum* secretes extracellular enzymes that facilitate cross-feeding of oligosaccharide degradation products among other *Bifidobacterium* species (Gotoh et al., 2018). Furthermore, *B. bifidum* strains are known to breakdown mucin which may aid gut colonization (Turroni et al., 2014). This personalized microbiota supplementation strategy, including genomic and phenotypic analysis of the probiotic strain, tailored to the nutritional environment of the receiving gut, is an important consideration for future studies.

Extremely low birth weight infants (<1000 g) represented the most vulnerable cohort in this study, and presented less abundance of genus *Bifidobacterium*, potentially due to several factors including; lengthened antibiotic courses (Ting et al., 2019), underdeveloped gut physiology (i.e. poorer gut motility, and thinner mucus layer (Neu, 2007)), and difficulties in establishing full enteral feeding (de Waard et al., 2019). Indeed, previous clinical studies have had difficulties evaluating the beneficial effects of supplementation in this at-risk cohort of preterm infants (Aceti et al., 2015; Escribano et al., 2018). Notably, although extremely low birth weight Bif/Lacto infants had lower *Bifidobacterium* abundance, supplementation in our study did enhance levels when compared to Control infants. Thus, from an intervention strategy perspective, daily and prolonged supplementation may contribute to faster (re)establishment of *Bifidobacterium*, which may also promote colonization resistance against exogenous or resident pathogens in this particularly fragile preterm cohort.

Surprisingly, *Bifidobacterium* abundance in supplemented infants was not affected by delivery mode (i.e. vaginal vs. caesarean-section), which has recently been shown in a large UK cohort of full-term infants (Shao et al., 2019). The early and frequent antibiotic treatment in preterm infants may create a ‘naïve niche’, which enables supplemented bifidobacterial strains to colonize successfully. Indeed, routine oral supplementation in Bif/Lacto infants may help to ameliorate caesarean delivery-associated bacterial alterations in the preterm microbiota and associated immune disorders (Goedicke-Fritz et al., 2017; Shao et al., 2019). Follow-up studies in this infant cohort could determine if this also contributes to reducing diseases associated with caesarean delivery or preterm birth such as asthma (Sonnenschein-van der Voort et al., 2014).

Rates of antibiotic prescription in preterms are remarkably high, ranging from 79% to 87% in extremely low birth (<1000 g) preterm infants (Bizzarro, 2018; Flannery et al., 2018). Antibiotic treatment favors the establishment of antibiotic-resistant bacteria, while indirectly eradicating highly susceptible microbiota members such as *Bifidobacterium* (Gasparrini et al., 2019; Rose et al., 2017). Recent research in infants correlated abundance of *Bifidobacterium* species (with/without supplementation) with a reduction in antimicrobial resistance genes and transferable elements (Esaiassen et al., 2018; Taft et al., 2018). As Bif/Lacto infants were predominantly colonized by genus *Bifidobacterium*, this may have contributed to reduce the reservoir of potentially multidrug resistance pathogens (i.e. *Staphylococcus, Escherichia*, and *Klebsiella*), which were prevalent in Control infants, and which have previously been shown to harbour a large repertoire of AMR determinants (including in this cohort) (Leggett RM, 2019; Rowe et al., 2019).

Previous studies have indicated that although preterm infants are particularly at risk of serious diseases with a bacterial cause (e.g. NEC and late onset sepsis), probiotic supplementation can reduce incidence (AlFaleh and Anabrees, 2014; Dermyshi et al., 2017). However, there has been variability in results, which may relate to the differences in strain(s) chosen, infant diet, or infant age. Notably, a recent clinical audit in the same NICU where the oral supplementation was given (i.e. Norfolk and Norwich University Hospital), indicated a >50% reduction in NEC rates and late-onset sepsis when comparing 5-year epochs before and after introducing probiotic supplementation, with no episodes of probiotic ‘sepsis’ indicated (P, 2019). Whilst the processes leading to life-threatening conditions including NEC in preterm infants are complex, overgrowth of potentially pathogenic bacteria is thought to be a key factor (Cassir et al., 2016; Leggett RM, 2019; Sim et al., 2015). We show that supplemented preterm infants have lower relative abundance of pathobionts including *Klebsiella* and *Escherichia*, which have previously been linked to NEC and late onset sepsis, and links to a recent study that performed MinION shotgun metagenomics and AMR profiling on samples from this cohort (Leggett RM, 2019). This may be due to direct inhibition through compounds secreted by *Bifidobacterium* (e.g. bacteriocins (Martinez et al., 2013)), competition for space and/or nutrients availability.

Limitations of this study include that it is observational in nature. However, low bifidobacteria abundance has been consistently reported in preterm infants in NICUs without any supplementation use (Alcon-Giner et al., 2017; Gasparrini et al., 2019; Sim et al., 2015). This indicates that the primary finding of high proportions of bifidobacteria, and associated changes in gut metabolites, in supplemented infants in this study is unlikely to be due to chance. Higher proportions of infants receiving probiotic supplements did receive mothers breast milk compared to a mix of breast milk and donor breast milk in the control. However, this did not result in any measurable difference in *Bifidobacterium* abundance or overall difference in microbiota composition. The HMOs in breast milk remain unaffected by the pasteurization and storage enabling donor milk to provide an equivalent substrate for the growth of bifidobacteria as breast milk (Bertino et al., 2008). Concerns have been raised about the safety of using probiotic bacteria in vulnerable individuals (Didari et al., 2014). However, no adverse effects were observed to result from over five years of routine clinical use of probiotics used in this study (P, 2019).

Differences in the gut environment were also indicated through our metabolomic analyses; highlighted by elevated abundance of acetate and lactate in feces from the Bif/Lacto Group, which are known to be primary metabolic end products of HMO degradation by *Bifidobacterium* (Kirmiz et al., 2018; Pokusaeva et al., 2011). Acetate and lactate have beneficial health effects enhancing defense functions in both host epithelial cells (Fukuda et al., 2012) and mucosal dendritic cells (Morita et al., 2019). The lower fecal pH in Bif/Lacto Group correlated with higher concentrations of these acids and higher abundance of *Bifidobacterium*, creating an acidic environment less favorable for the growth of pathobionts (Fukuda et al., 2011; Henrick et al., 2018; O’Sullivan et al., 2015).

In summary, we have provided a comprehensive basis for the beneficial impact of Bif/Lacto supplementation on the wider microbiota over time, providing a more mechanism-orientated approach to our data analysis. Although previous studies have investigated aspects of this before (Abdulkadir et al., 2016; Esaiassen et al., 2018; Plummer et al., 2018; Watkins et al., 2019), this is the largest study to combine multiple factors; fecal microbiota composition analysis, metabolomics, fecal pH, whole genome sequencing of supplemented probiotic strains and fecal isolates to confirm probiotic survival and colonisation, complemented by phenotypic testing. A key strength relates to the size and scope of the study; representing one of the largest reported longitudinal studies in preterm infants, where study cohorts were matched by gestational age, sex, birth-mode, time points of sample collection, and diet, which are all factors that may significantly impact the microbiota, and thus conclusions obtained. Alongside the key microbiological findings of this study, we have also provided context for further trials focusing on clinical practice in NICU and suggestions for future intervention studies in this at-risk infant population. Providing maternal and donor breast milk are essential practices required for successful gut colonization by *Bifidobacterium*, which also contributes to the enhanced metabolic end-products such as acetate and lactate in the preterm gut. These products will play an important role in direct antagonism of potentially pathogenic microbes, and the maturation of immune cells in early life. This large-scale longitudinal observational multi-center-controlled study emphasizes the important role that targeted microbiota or probiotic supplementation plays in preterm infants, exerting protective and functional effects on preterm gut microbial communities.

## Acknowledgements

This work was funded via a Wellcome Trust Investigator Award to LJH (100974/Z/13/Z) and support of the BBSRC Norwich Research Park Bioscience Doctoral Training Grant (BB/M011216/1, supervisor LJH, student CAG), Institute Strategic Programme (ISP) grant for Gut Health and Food Safety, BB/J004529/1 (LJH), and ISP grant for Gut Microbes and Health BB/R012490/1 and its constituent project(s), BBS/E/F/000PR10353 and BBS/E/F/000PR10355 to LJH. Work at St Mary’s was supported by a programme grant from the Winnicott Foundation to JSK, and the National Institute for Health Research (NIHR) Biomedical Research Centre based at Imperial Healthcare NHS Trust and Imperial College London. KS was funded by an NIHR Doctoral Research Fellowship [NIHR-DRF-2011-04-128]. The funders had no role in study design, data collection and analysis, decision to publish, or preparation of the manuscript. We sincerely thank all clinical nurses at Norfolk and Norwich University Hospital (NNUH), Rosie Hospital, Queen Charlotte’s and Chelsea Hospital, and St Mary’s Hospital for collecting stool samples. We would like to give a special mention to research nurses Karen Few, Hayley Aylmer and Zoe McClure for obtaining consent from parents and collecting samples. We thank Wellcome Trust Sanger Institute for their sequencing support.

## Author’s contributions

Overall study design was conceived by LJH, with PC, JSK, MJD and CAG contributing to refinement. Sample preparation for sequencing was performed by CAG, JK, CL, LC, MK, KLL, AS and KS. Bioinformatics and related computational analyses were performed by SC, RK and SM. Metabolomic analysis was performed by CAG, FFR and JRS. Phenotypic assays were performed by CAG and ML. Statistical analyses and final figure preparation was performed by MJD with input from LJH, CAG, SC, RK, ML, and JRS. LJH, CAG, and MJD wrote the manuscript with important contributions to intellectual content from all authors.

## Online Methods

### Study design

Two distinct preterm groups were recruited for this study: 1) Bif/Lacto Group who routinely received oral *Bifidobacterium* and *Lactobacillus* supplementation (n = 101 infants), and 2) Control Group infants who did not receive supplementation (n = 133 infants). Infants in the Bif/Lacto group were prescribed daily oral supplementation of 10^9^ colony forming units (CFU) of *Bifidobacterium bifidum* and 10^9^ CFU of *Lactobacillus acidophilus* (Infloran®, Desma Healthcare, Chiasso, Switzerland). This supplementation was given twice daily in a divided dose and commenced with the first enteral colostrum/milk feed (usually day 1 postnatal). Oral supplementation was normally administered until 34 weeks post-conceptual age, with the exception of very low birth weight infants (<1500 g) who received it until discharge. Half a capsule of Infloran (125 mg) was dissolved in 1 ml of expressed breastmilk and/or sterile water, and this dose was given twice daily (250mg/total/day) to the infant via nasogastric tube. Recruitment inclusion criteria included gestational age ≤ 34 weeks, and infants resident in the same NICU for study duration. Infants diagnosed with advanced stages of necrotizing enterocolitis or severe congenital abnormalities, were excluded from the study.

Preterm infants were recruited from four different NICUs across England, UK (between 2013-2017); Norfolk and Norwich University Hospital (NNUH) enrolled the Bif/Lacto group, and Rosie Hospital, Queen Charlotte’s and Chelsea Hospital, and St Mary’s Hospital recruited Control group infants. All NICUs had comparable health care practices including antibiotic and antifungal policies. Bif/Lacto vs. Control groups included similar sex ratios, delivery mode (i.e. Caesarean-section or vaginal delivery) and feeding modes (Supp Table 1). Specifically, we controlled for diet by preferably selecting preterm infants who received their mother’s own breast milk or donor breast milk; majority were exclusively breastfed or received donor breast milk (78% in Bif/Lacto Group and 76% Control Group), mixed fed with a combination of breastmilk, formula or donor breast milk (20% Bif/Lacto Group and 20% Control Group), and exclusively formula fed (2% Bif/Lacto Group, and 4% Control Group).

### Ethical approval for the study

Fecal collection from NNUH and Rosie Hospital was approved by the Faculty of Medical and Health Sciences Ethics Committee at the University of East Anglia (UEA), and followed protocols laid out by the UEA Biorepository (License no: 11208). Fecal collection for Queen Charlotte’s and Chelsea Hospital and St Mary’s Hospital was approved by West London Research Ethics Committee (REC) under the REC approval reference number 10/H0711/39. In all cases, doctors and nurses recruited infants after parents gave written consent.

Time points of sample collection for this study included 0-9 days, 10-29 days, 30-49 days, 50-99 days. Clinical data collected included gestational age, delivery mode, antibiotic courses received, and dietary information can be found in Supplementary Table 3.

### DNA extraction of preterm stool samples

FastDNA Spin Kit for Soil (MP) was used to extract DNA from preterm feces following manufacturer instructions, with extended 3 min bead-beating. DNA concentration and quality were quantified using a Qubit^®^ 2.0 fluorometer (Invitrogen).

### 16S rRNA gene sequencing: library preparation and bioinformatics analysis

16S rRNA region (V1-V2) primers were used for library construction (Supp Table 4), with the following PCR conditions; cycle of 94°C 3 min and 25 cycles of 94°C for 45 s, 55°C for 15 s and 72°C for 30 s. Sequencing of the 16S rRNA gene libraries was performed using Illumina MiSeq platform with 300 bp paired end reads.

Raw reads were filtered through quality control using trim galore (version 0.4.3), minimum quality threshold of phred 33, and minimum read length of 60 bp. Reads that passed threshold were aligned against SILVA database (version: SILVA_132_SSURef_tax_silva) using BLASTN (ncbi-blast-2.2.25+; Max e-value 10e-3) separately for both pairs. After performing BLASTN alignment, all output files were imported and annotated using the paired-end protocol of MEGAN on default Lowest Common Ancestor (LCA) parameters.

R Studio version 1.1.463 and using the ggplot2 R package version 3.1.0 was used for the analysis of microbiota sequence data and generation of figures. The 16S rRNA bacterial sequence data was subsampled to an even depth of 20,000 read using phyloseq package version 1.24.2. Sample details with proportion of reads assigned to each bacterial genus can be found in Supplementary Table 6 and species in Supplementary Table 7. NMDS (Non-metric multidimensional scaling) plots were generated with a Bray-Curtis dissimilarity calculation in R Studio using with the vegan package version 2.5-4 using code adapted from Torondel et al. (2016) (Torondel et al., 2016). Permutational MANOVA in the Adonis function of the vegan R package version 2.5-4 was used to determine significant differences between NMDS community structure. Heatmaps were generated using the ComplexHeatmap package version 1.18.1 and clustered using a Bray-Curtis dissimilarity calculation. Genus number, Shannon diversity, and Inverse Simpson diversity were calculated using the vegan package version 2.5-4. Statistically significant differences in genus and species abundance were determined using a Kruskal-Wallis test corrected for false discovery rate (FDR < 0.05). R scripts are available at: https://github.com/dalbymj/BAMBI-Paper-Files.

### Genomic DNA Extraction from bacterial isolates

We isolated the strains present in the oral supplementation (i.e. *Bifidobacterium bifidum* and *Lactobacillus acidophilus)* as well as additional *Bifidobacterium* isolates from infant samples. Overnight pure cultures in Brain Heart Infusion Broth (BHI) were harvested for phenol-chloroform DNA extraction. Bacterial pellets were resuspended in 2 ml 25% sucrose in 10 mM Tris and 1 mM EDTA at pH 8. Cells were subsequently lysed adding 50 μl 100 mg/ml lysozyme (Roche) and incubating at 37 °C for 1 h. 100 μl 20 mg/ml Proteinase K (Roche), 30 μl 10 mg/ml RNase A (Roche), 400 μl 0.5 M EDTA (pH 8.0) and 250 μl 10% Sarkosyl NL30 (Fisher) was added into the lysed bacterial suspension, incubated 1 h on ice and left overnight at 50 °C. Next, washes of phenol-chloroform-isoamyl alcohol (PCIA, Sigma) using 15 ml gel-lock tubes (Qiagen), with E Buffer (10mM Tris pH 8 (Fisher Scientific, UK)) added to sample to a final volume of 5 ml, mixed with 5 ml of PCIA (Sigma) and centrifuged for 15 min at 4000 rpm. The CIA step was repeated three times, after which the final aqueous phase was transferred into sterile Corning ^™^50 ml centrifuge tubes, and 2.5 volumes of ethanol (VWR Chemicals, USA) added, incubated for 15 min at - 20 °C, and centrifuged 10 min at 4000 rpm and 4 °C. Finally, the pellet was washed twice with 10 ml of 70% ethanol and centrifuged at 4000 rpm for 10 min, dried overnight, and re-suspended in 300 µl of E Buffer.

### Whole genome sequencing analysis: library preparation and bioinformatics analysis

DNA from pure cultures was sequenced at Wellcome Trust Sanger Institute using 96-plex Illumina HiSeq 2500 platform to generate 125 bp paired end reads as described previously (Harris et al., 2010). Genome assembly was performed by the sequencing provider using the assembly pipeline described by Page et al. 2016 (Page et al., 2016). Next, genome assemblies were annotated using Prokka (version 1.12). We predicted the 16s rRNA gene from the whole genome data using barrnap (version 0.7) and compare it to with existing 16S rRNA sequences. Single nucleotide variants (SNPs) were identified using Snippy (version 4.0) by mapping assembled contigs to annotated reference Infloran *B. bifidum* strain to reconstruct SNP phylogeny of six *B. bifidum* strains (Seemann, 2015).

To construct a phylogeny of 10 *Bifidobacterium* strains, we used pangenome pipeline Roary (version 3.12.0) to build a core gene alignment (87 core genes, with options -e -n otherwise default), followed by snp-sites (version 2.3.3) to call SNPs (6,202 SNPs in total) (Page et al., 2015; Page et al., 2016). We used the SNP site-alignments obtained from both reference-based and core-gene alignment approaches to infer Maximum Likelihood (ML) phylogenies using RAxML (version 8.2.10) with GTR+ nucleotide substitution model at 100 permutations conducted for bootstrap convergence test (Stamatakis, 2014). The ML tree reconstructed was with the highest likelihood out of 5 runs (option -N 5). Pairwise SNP distances were calculated and compared using snp-dists (version 0.2) (Seemann, 2018). Pairwise Average Nucleotide Identity (ANI) was computed and graphed using module pyani (version 0.2.7) (Pritchard et al., 2016). Web tool iTOL was used to visualize and annotate ML trees (Letunic and Bork, 2016).

### Determination of Minimal Inhibitory Concentration (MIC) of *Bifidobacterium bifidum* Infloran strain

The microdilution method was used to test Minimal Inhibitory Concentration (MIC) of the probiotic strains (*B. bifidum*) against routinely prescribed antibiotics; benzylpenicillin, gentamicin, and meropenem. Serial twofold dilutions of antibiotics in MRS medium (Difco) and 10 μL from fresh overnight culture were incubated for 24 h at 37°C under anaerobic conditions. Cell density was monitored using a plate reader (BMG Labtech, UK) at 595 nm. MICs were determined as the lowest concentration of antibiotic inhibiting any bacterial growth, with tests performed in triplicate.

### Gene search using BLAST

Genomes from *B. bifidum* Infloran strain and five other *B. bifidum* isolates were searched for genes involved in utilization of human milk oligosaccharides, and mucin degradation. Nucleotide sequences of genes of interest were extracted from National Centre of Biotechnology Information (NCBI).

Supplementary table 5 summarizes genes analyzed and publication source. BLAST alignment (ncbi-blast-2.2.25) was performed using a filtering criteria of 80% coverage and 80% identity.

### Breast milk and human milk oligosaccharides utilization study

Growth kinetics of the Infloran isolate *B. bifidum* and control type strains *B. longum* subsp. *infantis* DSM 20088 and *B. breve* DSM 20213 in breast milk and individual HMOs (LNnT or 2’FL) were performed. Isolates were grown overnight in RCM (Oxoid) then subcultured into modified MRS (Difco) with breast milk (pooled from multiple mothers, 1% w/v), or individual HMOs (2% w/v). Growth kinetics were measured every 15 minutes for 48 hours using a microplate spectrophotometer (Tecan Infinite F50).

### Metabolomic profiling of preterm feces using one dimensional 1D-nuclear magnetic resonance spectroscopy (NMR) and 2Dnuclear magnetic resonance spectroscopy (NMR)

A subset of 157 paired fecal samples (75 from Bif/Lacto group, and 81 from Control group) were analyzed by standard one-dimensional (1D) ^1^H NMR spectroscopy using a Bruker 600 MHz spectrometer operating at 300 K. Fecal samples were chosen for metabolic profiling pragmatically based on remaining sample quantity after previous analyses. Feces (50 mg) were combined with 700 μl of phosphate buffer (pH 7.4; 100% D_2_O) containing 1 mmol/L of 3-trimethylsilyl-1-[2,2,3,3-^2^H_4_] propionate (TSP), and 10 zirconium beads (1 mm diameter) (BioSpec Products). Samples were homogenized using a Precellys bead beater (Bertin) with 2 cycles of 40 s at 6,500 Hz speed, centrifuged at 14,000 *g* for 10 min and the supernatant was transferred to NMR tubes. 1D NMR spectra were acquired for each sample using a nuclear overhauser effect pulse sequence for water suppression as described by Beckonert and colleagues (Beckonert et al., 2007)). Spectra were automatically phased and calibrated to the TSP reference using Topspin 3.6 (Bruker BioSpin). Spectra were imported into Matlab 9.4 (R2018a), redundant spectral regions (those arising from TSP and imperfect water suppression) were removed, and the spectral profiles were normalized using a probabilistic quotient method. Data analysis was performed using principal components analysis (PCA), orthogonal projection to latent structures discriminant analysis (OPLS-DA) and covariate-adjusted-projection to latent structures-discriminant analysis (CA-PLSDA) using in-house scripts. Pairwise OPLS-DA models (Bif/Lacto versus Control) were constructed for each sampling point and for all sampling points combined. Here, the complete spectral data points (metabolic profile) served as the predictors (X variables) and class membership (Bif/Lacto versus Control) served as the response (Y) variable. The predictive ability (Q^2^Y) of the models were calculated using a 7-fold cross-validation approach and the validity of the Q^2^Y values were assessed through permutation testing (100 permutations). A CA-PLS model was also built using the fecal profiles from all sampling points and the model was adjusted for sampling age. Additional two-dimensional (2D) ^1^H-^1^H NMR spectroscopy was performed on two selected fecal samples to assist with metabolite identification. Conventional 2D NMR spectra were acquired using homonuclear correlation spectroscopy (COSY) and heteronuclear single quantum coherence spectroscopy (HSQC) experiments with water suppression to assist with structural elucidation.

### pH measurement of the fecal samples

The pH of a randomly selected subset of fecal samples used in the metabolomics analysis (39 samples from the Bif/Lacto Group, and 39 samples from the Control Group) was assessed. Fifty mg of fecal sample was added 1 ml of sterile water, vortexed and measured using a glass electrode pH meter (Martini Mi151).

### Accession codes

The datasets supporting the conclusions of this article are available in the European Nucleotide Archive (http://www.ebi.ac.uk/ena repository) accession numbers PRJEB31653 (Illumina 16S MiSeq sequencing and Illumina HiSeq sequencing data).

## References

Abdulkadir, B., Nelson, A., Skeath, T., Marrs, E.C., Perry, J.D., Cummings, S.P., Embleton, N.D., Berrington, J.E., and Stewart, C.J. (2016). Routine Use of Probiotics in Preterm Infants: Longitudinal Impact on the Microbiome and Metabolome. Neonatology. 109(4), 239–247. Published online 2016/02/10 DOI: 10.1159/000442936.

Aceti, A., Gori, D., Barone, G., Callegari, M.L., Di Mauro, A., Fantini, M.P., Indrio, F., Maggio, L., Meneghin, F., Morelli, L., et al. (2015). Probiotics for prevention of necrotizing enterocolitis in preterm infants: systematic review and meta-analysis. Ital J Pediatr. 41, 89. Published online 2015/11/17 DOI: 10.1186/s13052-015-0199-2.

Alcon-Giner, C., Caim, S., Mitra, S., Ketskemety, J., Wegmann, U., Wain, J., Belteki, G., Clarke, P., and Hall, L.J. (2017). Optimisation of 16S rRNA gut microbiota profiling of extremely low birth weight infants. BMC Genomics. 18(1), 841. Published online 2017/11/04 DOI: 10.1186/s12864-017-4229-x.

AlFaleh, K., and Anabrees, J. (2014). Probiotics for prevention of necrotizing enterocolitis in preterm infants. Cochrane Database Syst Rev. (4), CD005496. Published online 2014/04/12 DOI: 10.1002/14651858.CD005496.pub4.

Ayechu-Muruzabal, V., van Stigt, A.H., Mank, M., Willemsen, L.E.M., Stahl, B., Garssen, J., and Van’t Land, B. (2018). Diversity of Human Milk Oligosaccharides and Effects on Early Life Immune Development. Front Pediatr. 6, 239. Published online 2018/09/27 DOI: 10.3389/fped.2018.00239.

Beckonert, O., Keun, H.C., Ebbels, T.M., Bundy, J., Holmes, E., Lindon, J.C., and Nicholson, J.K. (2007). Metabolic profiling, metabolomic and metabonomic procedures for NMR spectroscopy of urine, plasma, serum and tissue extracts. Nat Protoc. 2(11), 2692–2703. Published online 2007/11/17 DOI: 10.1038/nprot.2007.376.

Been, J.V., Lugtenberg, M.J., Smets, E., van Schayck, C.P., Kramer, B.W., Mommers, M., and Sheikh, A. (2014). Preterm birth and childhood wheezing disorders: a systematic review and meta-analysis. PLoS Med. 11(1), e1001596. Published online 2014/02/05 DOI: 10.1371/journal.pmed.1001596.

Bertino, E., Coppa, G.V., Giuliani, F., Coscia, A., Gabrielli, O., Sabatino, G., Sgarrella, M., Testa, T., Zampini, L., and Fabris, C. (2008). Effects of Holder pasteurization on human milk oligosaccharides. Int J Immunopathol Pharmacol. 21(2), 381–385. Published online 2008/06/13 DOI: 10.1177/039463200802100216.

Bizzarro, M.J. (2018). Avoiding Unnecessary Antibiotic Exposure in Premature Infants: Understanding When (Not) to Start and When to Stop. JAMA Netw Open. 1(1), e180165. Published online 2019/01/16 DOI: 10.1001/jamanetworkopen.2018.0165.

Cassir, N., Simeoni, U., and La Scola, B. (2016). Gut microbiota and the pathogenesis of necrotizing enterocolitis in preterm neonates. Future Microbiol. 11(2), 273–292. Published online 2016/02/09 DOI: 10.2217/fmb.15.136.

Costeloe, K., Hardy, P., Juszczak, E., Wilks, M., Millar, M.R., and Study, P.P.I. (2016). Bifidobacterium breve BBG-001 in very preterm infants: a randomised controlled phase 3 trial. Lancet. 387(10019), 649–660.

Dahl, C., Stigum, H., Valeur, J., Iszatt, N., Lenters, V., Peddada, S., Bjornholt, J.V., Midtvedt, T., Mandal, S., and Eggesbo, M. (2018). Preterm infants have distinct microbiomes not explained by mode of delivery, breastfeeding duration or antibiotic exposure. Int J Epidemiol. 47(5), 1658–1669. Published online 2018/04/25 DOI: 10.1093/ije/dyy064.

de Waard, M., Li, Y., Zhu, Y., Ayede, A.I., Berrington, J., Bloomfield, F.H., Busari, O.O., Cormack, B.E., Embleton, N.D., van Goudoever, J.B., et al. (2019). Time to Full Enteral Feeding for Very Low-Birth-Weight Infants Varies Markedly Among Hospitals Worldwide But May Not Be Associated With Incidence of Necrotizing Enterocolitis: The NEOMUNE-NeoNutriNet Cohort Study. JPEN J Parenter Enteral Nutr. 43(5), 658–667. Published online 2018/11/23 DOI: 10.1002/jpen.1466.

Dermyshi, E., Wang, Y., Yan, C., Hong, W., Qiu, G., Gong, X., and Zhang, T. (2017). The “Golden Age” of Probiotics: A Systematic Review and Meta-Analysis of Randomized and Observational Studies in Preterm Infants. Neonatology. 112(1), 9–23. Published online 2017/02/15 DOI: 10.1159/000454668.

Deshpande, G., Rao, S., Athalye-Jape, G., Conway, P., and Patole, S. (2016). Probiotics in very preterm infants: the PiPS trial. Lancet. 388(10045), 655–655.

Didari, T., Solki, S., Mozaffari, S., Nikfar, S., and Abdollahi, M. (2014). A systematic review of the safety of probiotics. Expert Opin Drug Saf. 13(2), 227–239. Published online 2014/01/11 DOI: 10.1517/14740338.2014.872627.

Dominguez-Bello, M.G., Costello, E.K., Contreras, M., Magris, M., Hidalgo, G., Fierer, N., and Knight, R. (2010). Delivery mode shapes the acquisition and structure of the initial microbiota across multiple body habitats in newborns. Proc Natl Acad Sci U S A. 107(26), 11971–11975. Published online 2010/06/23 DOI: 10.1073/pnas.1002601107.

Duffield, S.D., and Clarke, P. (2019). Current use of probiotics to prevent necrotising enterocolitis. Arch Dis Child Fetal Neonatal Ed. 104(2), F228. Published online 2018/11/23 DOI: 10.1136/archdischild-2018-316199.

Esaiassen, E., Hjerde, E., Cavanagh, J.P., Pedersen, T., Andresen, J.H., Rettedal, S.I., Stoen, R., Nakstad, B., Willassen, N.P., and Klingenberg, C. (2018). Effects of Probiotic Supplementation on the Gut Microbiota and Antibiotic Resistome Development in Preterm Infants. Front Pediatr. 6, 347. Published online 2018/12/07 DOI: 10.3389/fped.2018.00347.

Escribano, E., Zozaya, C., Madero, R., Sanchez, L., van Goudoever, J., Rodriguez, J.M., and de Pipaon, M.S. (2018). Increased incidence of necrotizing enterocolitis associated with routine administration of Infloran in extremely preterm infants. Benef Microbes. 9(5), 683–690. Published online 2018/06/12 DOI: 10.3920/BM2017.0098.

Flannery, D.D., Ross, R.K., Mukhopadhyay, S., Tribble, A.C., Puopolo, K.M., and Gerber, J.S. (2018). Temporal Trends and Center Variation in Early Antibiotic Use Among Premature Infants. JAMA Netw Open. 1(1), e180164. Published online 2019/01/16 DOI: 10.1001/jamanetworkopen.2018.0164.

Fukuda, S., Toh, H., Hase, K., Oshima, K., Nakanishi, Y., Yoshimura, K., Tobe, T., Clarke, J.M., Topping, D.L., Suzuki, T., et al. (2011). Bifidobacteria can protect from enteropathogenic infection through production of acetate. Nature. 469(7331), 543–547. Published online 2011/01/29 DOI: 10.1038/nature09646.

Fukuda, S., Toh, H., Taylor, T.D., Ohno, H., and Hattori, M. (2012). Acetate-producing bifidobacteria protect the host from enteropathogenic infection via carbohydrate transporters. Gut Microbes. 3(5), 449–454. Published online 2012/07/25 DOI: 10.4161/gmic.21214.

Gasparrini, A.J., Wang, B., Sun, X., Kennedy, E.A., Hernandez-Leyva, A., Ndao, I.M., Tarr, P.I., Warner, B.B., and Dantas, G. (2019). Persistent metagenomic signatures of early-life hospitalization and antibiotic treatment in the infant gut microbiota and resistome. Nat Microbiol. Published online 2019/09/11 DOI: 10.1038/s41564-019-0550-2.

Goedicke-Fritz, S., Hartel, C., Krasteva-Christ, G., Kopp, M.V., Meyer, S., and Zemlin, M. (2017). Preterm Birth Affects the Risk of Developing Immune-Mediated Diseases. Front Immunol. 8, 1266. Published online 2017/10/25 DOI: 10.3389/fimmu.2017.01266.

Gotoh, A., Katoh, T., Sakanaka, M., Ling, Y., Yamada, C., Asakuma, S., Urashima, T., Tomabechi, Y., Katayama-Ikegami, A., Kurihara, S., et al. (2018). Sharing of human milk oligosaccharides degradants within bifidobacterial communities in faecal cultures supplemented with Bifidobacterium bifidum. Sci Rep. 8(1), 13958. Published online 2018/09/20 DOI: 10.1038/s41598-018-32080-3.

Haataja, P., Korhonen, P., Ojala, R., Hirvonen, M., Paassilta, M., Gissler, M., Luukkaala, T., and Tammela, O. (2016). Asthma and atopic dermatitis in children born moderately and late preterm. Eur J Pediatr. 175(6), 799–808. Published online 2016/02/24 DOI: 10.1007/s00431-016-2708-8.

Harris, S.R., Feil, E.J., Holden, M.T., Quail, M.A., Nickerson, E.K., Chantratita, N., Gardete, S., Tavares, A., Day, N., Lindsay, J.A., et al. (2010). Evolution of MRSA during hospital transmission and intercontinental spread. Science. 327(5964), 469–474. Published online 2010/01/23 DOI: 10.1126/science.1182395.

Henderickx, J.G.E., Zwittink, R.D., van Lingen, R.A., Knol, J., and Belzer, C. (2019). The Preterm Gut Microbiota: An Inconspicuous Challenge in Nutritional Neonatal Care. Front Cell Infect Microbiol. 9, 85. Published online 2019/04/20 DOI: 10.3389/fcimb.2019.00085.

Henrick, B.M., Hutton, A.A., Palumbo, M.C., Casaburi, G., Mitchell, R.D., Underwood, M.A., Smilowitz, J.T., and Frese, S.A. (2018). Elevated Fecal pH Indicates a Profound Change in the Breastfed Infant Gut Microbiome Due to Reduction of Bifidobacterium over the Past Century. mSphere. 3(2). Published online 2018/03/23 DOI: 10.1128/mSphere.00041-18.

Hickey, L., Garland, S.M., Jacobs, S.E., O’Donnell, C.P., Tabrizi, S.N., and ProPrems Study, G. (2014). Cross-colonization of infants with probiotic organisms in a neonatal unit. J Hosp Infect. 88(4), 226–229. Published online 2014/12/03 DOI: 10.1016/j.jhin.2014.09.006.

Hill, C., Guarner, F., Reid, G., Gibson, G.R., Merenstein, D.J., Pot, B., Morelli, L., Canani, R.B., Flint, H.J., Salminen, S., et al. (2014). Expert consensus document. The International Scientific Association for Probiotics and Prebiotics consensus statement on the scope and appropriate use of the term probiotic. Nat Rev Gastroenterol Hepatol. 11(8), 506–514. Published online 2014/06/11 DOI: 10.1038/nrgastro.2014.66.

Kirmiz, N., Robinson, R.C., Shah, I.M., Barile, D., and Mills, D.A. (2018). Milk Glycans and Their Interaction with the Infant-Gut Microbiota. Annu Rev Food Sci Technol. 9, 429–450. Published online 2018/03/28 DOI: 10.1146/annurev-food-030216-030207.

Korpela, K., Blakstad, E.W., Moltu, S.J., Strommen, K., Nakstad, B., Ronnestad, A.E., Braekke, K., Iversen, P.O., Drevon, C.A., and de Vos, W. (2018). Intestinal microbiota development and gestational age in preterm neonates. Sci Rep. 8(1), 2453. Published online 2018/02/08 DOI: 10.1038/s41598-018-20827-x.

Leggett RM, A.-G.C., Heavens D, Caim S, Brook TC, Kujawska M, Martin S, Peel N, Acford-Palmer H, Hoyles4 L, Clarke P, Hall LJ, Clark MD (2019). Rapid MinION profiling of preterm microbiota and antimicrobial resistant pathogens. Nature Microbiology. Accepted

Letunic, I., and Bork, P. (2016). Interactive tree of life (iTOL) v3: an online tool for the display and annotation of phylogenetic and other trees. Nucleic Acids Res. 44(W1), W242–245. Published online 2016/04/21 DOI: 10.1093/nar/gkw290.

Martinez, F.A., Balciunas, E.M., Converti, A., Cotter, P.D., and de Souza Oliveira, R.P. (2013). Bacteriocin production by Bifidobacterium spp. A review. Biotechnol Adv. 31(4), 482–488. Published online 2013/02/07 DOI: 10.1016/j.biotechadv.2013.01.010.

Milani, C., Mancabelli, L., Lugli, G.A., Duranti, S., Turroni, F., Ferrario, C., Mangifesta, M., Viappiani, A., Ferretti, P., Gorfer, V., et al. (2015). Exploring Vertical Transmission of Bifidobacteria from Mother to Child. Appl Environ Microbiol. 81(20), 7078–7087. Published online 2015/08/02 DOI: 10.1128/AEM.02037-15.

Morita, N., Umemoto, E., Fujita, S., Hayashi, A., Kikuta, J., Kimura, I., Haneda, T., Imai, T., Inoue, A., Mimuro, H., et al. (2019). GPR31-dependent dendrite protrusion of intestinal CX3CR1(+) cells by bacterial metabolites. Nature. 566(7742), 110–114. Published online 2019/01/25 DOI: 10.1038/s41586-019-0884-1.

Mulder, I.E., Schmidt, B., Lewis, M., Delday, M., Stokes, C.R., Bailey, M., Aminov, R.I., Gill, B.P., Pluske, J.R., Mayer, C.D., et al. (2011). Restricting microbial exposure in early life negates the immune benefits associated with gut colonization in environments of high microbial diversity. PLoS One. 6(12), e28279. Published online 2012/01/05 DOI: 10.1371/journal.pone.0028279.

Neu, J. (2007). Gastrointestinal development and meeting the nutritional needs of premature infants. Am J Clin Nutr. 85(2), 629S–634S. Published online 2007/02/08 DOI: 10.1093/ajcn/85.2.629S.

O’Sullivan, A., Farver, M., and Smilowitz, J.T. (2015). The Influence of Early Infant-Feeding Practices on the Intestinal Microbiome and Body Composition in Infants. Nutr Metab Insights. 8(Suppl 1), 1–9. Published online 2015/12/31 DOI: 10.4137/NMI.S29530.

P, R.C.S.G.C.R.J.J.M.H.B.E.M.A.H.L.C. (2019). Incidence of necrotising enterocolitis before and after introducing routine prophylactic Lactobacillus and Bifidobacterium probiotics Archives of Disease in Childhood. In press.

Page, A.J., Cummins, C.A., Hunt, M., Wong, V.K., Reuter, S., Holden, M.T., Fookes, M., Falush, D., Keane, J.A., and Parkhill, J. (2015). Roary: rapid large-scale prokaryote pan genome analysis. Bioinformatics. 31(22), 3691–3693. Published online 2015/07/23 DOI: 10.1093/bioinformatics/btv421.

Page, A.J., Taylor, B., Delaney, A.J., Soares, J., Seemann, T., Keane, J.A., and Harris, S.R. (2016). SNP-sites: rapid efficient extraction of SNPs from multi-FASTA alignments. Microb Genom. 2(4), e000056. Published online 2017/03/30 DOI: 10.1099/mgen.0.000056.

Pammi, M., and Weisman, L.E. (2015). Late-onset sepsis in preterm infants: update on strategies for therapy and prevention. Expert Rev Anti Infect Ther. 13(4), 487–504. Published online 2015/02/11 DOI: 10.1586/14787210.2015.1008450.

Peisl, B.Y.L., Schymanski, E.L., and Wilmes, P. (2018). Dark matter in host-microbiome metabolomics: Tackling the unknowns-A review. Anal Chim Acta. 1037, 13–27. Published online 2018/10/08 DOI: 10.1016/j.aca.2017.12.034.

Plummer, E.L., Bulach, D.M., Murray, G.L., Jacobs, S.E., Tabrizi, S.N., Garland, S.M., and ProPrems Study, G. (2018). Gut microbiota of preterm infants supplemented with probiotics: sub-study of the ProPrems trial. BMC Microbiol. 18(1), 184. Published online 2018/11/15 DOI: 10.1186/s12866-018-1326-1.

Pokusaeva, K., Fitzgerald, G.F., and van Sinderen, D. (2011). Carbohydrate metabolism in Bifidobacteria. Genes Nutr. 6(3), 285–306. Published online 2011/04/13 DOI: 10.1007/s12263-010-0206-6.

Pritchard, L., Glover, R.H., Humphris, S., Elphinstone, J.G., and Toth, I.K. (2016). Genomics and taxonomy in diagnostics for food security: soft-rotting enterobacterial plant pathogens. Anal Methods-Uk. 8(1), 12–24.

Rose, G., Shaw, A.G., Sim, K., Wooldridge, D.J., Li, M.S., Gharbia, S., Misra, R., and Kroll, J.S. (2017). Antibiotic resistance potential of the healthy preterm infant gut microbiome. PeerJ. 5, e2928. Published online 2017/02/06 DOI: 10.7717/peerj.2928.

Rowe, W.P., Carrieri, A.P., Alcon-Giner, C., Caim, S., Shaw, A., Sim, K., Kroll, J.S., Hall, L.J., Pyzer-Knapp, E.O., and Winn, M.D. (2019). Streaming histogram sketching for rapid microbiome analytics. Microbiome. 7(1), 40. Published online 2019/03/18 DOI: 10.1186/s40168-019-0653-2.

Seemann, T., 2015. Snippy: Rapid haploid variant calling and core genome alignment. GitHub.

Seemann, T.K. F.; Page, A.J, 2018. Snp-dists: Pairwise SNP distance matrix from a FASTA sequence alignment. GitHub.

Shao, Y., Forster, S.C., Tsaliki, E., Vervier, K., Strang, A., Simpson, N., Kumar, N., Stares, M.D., Rodger, A., Brocklehurst, P., et al. (2019). Stunted microbiota and opportunistic pathogen colonization in caesarean-section birth. Nature. Published online 2019/09/20 DOI: 10.1038/s41586-019-1560-1.

Shulhan, J., Dicken, B., Hartling, L., and Larsen, B.M. (2017). Current Knowledge of Necrotizing Enterocolitis in Preterm Infants and the Impact of Different Types of Enteral Nutrition Products. Adv Nutr. 8(1), 80–91. Published online 2017/01/18 DOI: 10.3945/an.116.013193.

Sim, K., Shaw, A.G., Randell, P., Cox, M.J., McClure, Z.E., Li, M.S., Haddad, M., Langford, P.R., Cookson, W.O., Moffatt, M.F., et al. (2015). Dysbiosis anticipating necrotizing enterocolitis in very premature infants. Clin Infect Dis. 60(3), 389–397. Published online 2014/10/26 DOI: 10.1093/cid/ciu822.

Sonnenschein-van der Voort, A.M., Arends, L.R., de Jongste, J.C., Annesi-Maesano, I., Arshad, S.H., Barros, H., Basterrechea, M., Bisgaard, H., Chatzi, L., Corpeleijn, E., et al. (2014). Preterm birth, infant weight gain, and childhood asthma risk: a meta-analysis of 147,000 European children. J Allergy Clin Immunol. 133(5), 1317–1329. Published online 2014/02/18 DOI: 10.1016/j.jaci.2013.12.1082.

Stamatakis, A. (2014). RAxML version 8: a tool for phylogenetic analysis and post-analysis of large phylogenies. Bioinformatics. 30(9), 1312–1313. Published online 2014/01/24 DOI: 10.1093/bioinformatics/btu033.

Suez, J., Zmora, N., Segal, E., and Elinav, E. (2019). The pros, cons, and many unknowns of probiotics. Nat Med. 25(5), 716–729. Published online 2019/05/08 DOI: 10.1038/s41591-019-0439-x.

Taft, D.H., Liu, J., Maldonado-Gomez, M.X., Akre, S., Huda, M.N., Ahmad, S.M., Stephensen, C.B., and Mills, D.A. (2018). Bifidobacterial Dominance of the Gut in Early Life and Acquisition of Antimicrobial Resistance. mSphere. 3(5). Published online 2018/09/28 DOI: 10.1128/mSphere.00441-18.

Tauchi, H., Yahagi, K., Yamauchi, T., Hara, T., Yamaoka, R., Tsukuda, N., Watakabe, Y., Tajima, S., Ochi, F., Iwata, H., et al. (2019). Gut microbiota development of preterm infants hospitalised in intensive care units. Benef Microbes. 1–12. Published online 2019/06/11 DOI: 10.3920/BM2019.0003.

Timmerman, H.M., Rutten, N., Boekhorst, J., Saulnier, D.M., Kortman, G.A.M., Contractor, N., Kullen, M., Floris, E., Harmsen, H.J.M., Vlieger, A.M., et al. (2017). Intestinal colonisation patterns in breastfed and formula-fed infants during the first 12 weeks of life reveal sequential microbiota signatures. Sci Rep. 7(1), 8327. Published online 2017/08/23 DOI: 10.1038/s41598-017-08268-4.

Ting, J.Y., Roberts, A., Sherlock, R., Ojah, C., Cieslak, Z., Dunn, M., Barrington, K., Yoon, E.W., Shah, P.S., and Canadian Neonatal Network, I. (2019). Duration of Initial Empirical Antibiotic Therapy and Outcomes in Very Low Birth Weight Infants. Pediatrics. 143(3). Published online 2019/03/02 DOI: 10.1542/peds.2018-2286.

Torondel, B., Ensink, J.H., Gundogdu, O., Ijaz, U.Z., Parkhill, J., Abdelahi, F., Nguyen, V.A., Sudgen, S., Gibson, W., Walker, A.W., et al. (2016). Assessment of the influence of intrinsic environmental and geographical factors on the bacterial ecology of pit latrines. Microb Biotechnol. 9(2), 209–223. Published online 2016/02/16 DOI: 10.1111/1751-7915.12334.

Turroni, F., Duranti, S., Bottacini, F., Guglielmetti, S., Van Sinderen, D., and Ventura, M. (2014). Bifidobacterium bifidum as an example of a specialized human gut commensal. Front Microbiol. 5, 437. Published online 2014/09/06 DOI: 10.3389/fmicb.2014.00437.

Walker, W.A. (2017). The importance of appropriate initial bacterial colonization of the intestine in newborn, child, and adult health. Pediatr Res. 82(3), 387–395. Published online 2017/04/21 DOI: 10.1038/pr.2017.111.

Watkins, C., Murphy, K., Dempsey, E.M., O’Shea, C.A., Murphy, B.P., O’Toole, P.W., Ross, R.P., Stanton, C., and Ryan, C.A. (2019). Dose-interval study of a dual probiotic in preterm infants. Arch Dis Child Fetal Neonatal Ed. 104(2), F159–F164. Published online 2018/06/22 DOI: 10.1136/archdischild-2017-313468.

WHO, 2018. Preterm birth.

Zou, Z.H., Liu, D., Li, H.D., Zhu, D.P., He, Y., Hou, T., and Yu, J.L. (2018). Prenatal and postnatal antibiotic exposure influences the gut microbiota of preterm infants in neonatal intensive care units. Ann Clin Microbiol Antimicrob. 17(1), 9. Published online 2018/03/21 DOI: 10.1186/s12941-018-0264-y.

