## Supplementary Table 2 for "Microbiota supplementation with *Bifidobacterium* and *Lactobacillus* modifies the preterm infant gut microbiota and metabolome"

**Supplementary Table 2 -** AMR gene analysis and phenotypic characterisation of *Bifidobacterium bifidum* (Infloran)

**a** *Bifidobacterium bifidum* (Infloran)

| Gene | Identity (%) | Resistance to |
|---|---|---|
| **ileS** | 99.13 | Mupirocin |

**b**

| Antibiotic | ***Bifidobacterium bifidum*** MIC (mg/L) | Eucast value (mg/L) |
|---|---|---|
| Benzylpenicillin | 0.11 | 0.12 (ampicillin) |
| Gentamicin | 39 | 64 |
| Meropenem | 0.095 | ND |
