## Supplementary Table 4 for "Microbiota supplementation with *Bifidobacterium* and *Lactobacillus* modifies the preterm infant gut microbiota and metabolome"

*Supplementary Table 4* ***: Primer sequences for amplifying V1+V2 region of 16S rRNA gene using MiSeq Illumina.***

| 9 forward (FW) primers<br>12 reverse (RV) primers | Total Primer sequences |
|---|---|
| V1FW_SD501 | AATGATACGGCGACCACCGAGATCTACACAAGCAGCATATGGTAATTGTAGMGTTYGATYMTGGCTCAG |
| V1FW_SD502 | AATGATACGGCGACCACCGAGATCTACACACGCGTGATATGGTAATTGTAGMGTTYGATYMTGGCTCAG |
| V1FW_SD503 | AATGATACGGCGACCACCGAGATCTACACCGATCTACTATGGTAATTGTAGMGTTYGATYMTGGCTCAG |
| V1FW_SD504 | AATGATACGGCGACCACCGAGATCTACACTGCGTCACTATGGTAATTGTAGMGTTYGATYMTGGCTCAG |
| V1FW_SD505 | AATGATACGGCGACCACCGAGATCTACACGTCTAGTGTATGGTAATTGTAGMGTTYGATYMTGGCTCAG |
| V1FW_SD506 | AATGATACGGCGACCACCGAGATCTACACCTAGTATGTATGGTAATTGTAGMGTTYGATYMTGGCTCAG |
| V1FW_SD507 | AATGATACGGCGACCACCGAGATCTACACGATAGCGTTATGGTAATTGTAGMGTTYGATYMTGGCTCAG |
| V1FW_SD508 | AATGATACGGCGACCACCGAGATCTACACTCTACACTTATGGTAATTGTAGMGTTYGATYMTGGCTCAG |
| V1FW_SA501 | AATGATACGGCGACCACCGAGATCTACACATCGTACGTATGGTAATTGTAGMGTTYGATYMTGGCTCAG |
| V2RV_SD701 | CAAGCAGAAGACGGCATACGAGATACCTAGTAAGTCAGTCAGCCGCTGCCTCCCGTAGGAGT |
| V2RV_SD702 | CAAGCAGAAGACGGCATACGAGATACGTACGTAGTCAGTCAGCCGCTGCCTCCCGTAGGAGT |
| V2RV_SD703 | CAAGCAGAAGACGGCATACGAGATATATCGCGAGTCAGTCAGCCGCTGCCTCCCGTAGGAGT |
| V2RV_SD704 | CAAGCAGAAGACGGCATACGAGATCACGATAGAGTCAGTCAGCCGCTGCCTCCCGTAGGAGT |
| V2RV_SD705 | CAAGCAGAAGACGGCATACGAGATCGTATCGCAGTCAGTCAGCCGCTGCCTCCCGTAGGAGT |
| V2RV_SD706 | CAAGCAGAAGACGGCATACGAGATCTGCGACTAGTCAGTCAGCCGCTGCCTCCCGTAGGAGT |
| V2RV_SD707 | CAAGCAGAAGACGGCATACGAGATGCTGTAACAGTCAGTCAGCCGCTGCCTCCCGTAGGAGT |
| V2RV_SD708 | CAAGCAGAAGACGGCATACGAGATGGACGTTAAGTCAGTCAGCCGCTGCCTCCCGTAGGAGT |
| V2RV_SD709 | CAAGCAGAAGACGGCATACGAGATGGTCGTAGAGTCAGTCAGCCGCTGCCTCCCGTAGGAGT |
| V2RV_SD710 | CAAGCAGAAGACGGCATACGAGATTAAGTCTCAGTCAGTCAGCCGCTGCCTCCCGTAGGAGT |
| V2RV_SD711 | CAAGCAGAAGACGGCATACGAGATTACACAGTAGTCAGTCAGCCGCTGCCTCCCGTAGGAGT |
| V2RV_SD712 | CAAGCAGAAGACGGCATACGAGATTTGACGCAAGTCAGTCAGCCGCTGCCTCCCGTAGGAGT |
