## Supplementary Table 5 for "Microbiota supplementation with *Bifidobacterium* and *Lactobacillus* modifies the preterm infant gut microbiota and metabolome"

**Supplementary table 5 - *B.bifidum* HMOs and mucin degradation genes**

| Gene | Function | Publication |
|---|---|---|
| *lnbB* | HMO degradation | Biosidase, a Critical Enzyme for the Degradation of Human Milk Oligosaccharides with a Type 1 Structure. *Applied and Environmental Microbiology* **74**, 3996-4004 (2008). |
| *afcB, AfcA* | | Two distinct α-l-fucosidases from Bifidobacterium bifidum are essential for the utilization of fucosylated milk oligosaccharides and glycoconjugates. *Glycobiology* **19**, 1010-1017 (2009) |
| BbhI, BbgIII | | Cooperation of β-galactosidase and β-N-acetylhexosaminidase from bifidobacteria in assimilation of human milk oligosaccharides with type 2 structure. *Glycobiology* **20**, 1402-1409 (2010) |
| *Siabb2* | | Extracellular Sialidase Enhances Adhesion to the Mucosal Surface and Supports Carbohydrate Assimilation. *mBio* **8**, e00928-00917 (2017). |
| BBPR_0193, | Mucin degradation genes | Genome analysis of Bifidobacterium bifidum PRL2010 reveals metabolic pathways for host-derived glycan foraging. Proc Natl Acad Sci U S A 107, 19514-19519 (2010) |
| BBPR_1360 | | |
| BBPR_1793 | | |
| BBPR_0482 | | |
| BBPR_0264 | | |
| BBPR_1514 | | |
| BBPR_1018 | | |
