## Supplementary Figures for "Microbiota supplementation with *Bifidobacterium* and *Lactobacillus* modifies the preterm infant gut microbiota and metabolome"

**Supplementary Table 1: Summary statistics for preterm infants recruited in the study**

| | Group | | |
|---|---|---|---|
| | Control | Bif/Lacto | p-value |
| **n** | **133** | **101** | |
| **Sex (n (%))** | | | |
| *Female* | 75 (56.4) | 45 (44.6) | 0.096 |
| *Male* | 58 (43.6) | 56 (55.4) | |
| **Delivery (n (%))** | | | |
| *Vaginal* | 61 (45.9) | 46 (45.5) | 1 |
| *Cesarean* | 72 (54.1) | 55 (54.5) | |
| **Birthweight in grams (mean (SD))** | 1146.02 (337.19) | 1127.17 (311.97) | 0.662 |
| **Gestational age in weeks (mean (SD))** | 28.35 (2.29) | 28.55 (2.77) | 0.545 |
| **Length of NICU stay (days) (mean (SD))** | 45.86 (22.79) | 53.06 (32.71) | 0.055 |
| **Length of antibiotics (n (%))** | | | |
| *Long* | 25 (18.8) | 42 (41.6) | 0.001 |
| *Short* | 94 (70.7) | 54 (53.5) | |
| *None* | 13 (9.8) | 5 (5.0) | |
| *NA* | 1 (0.8) | 0 (0.0) | |
| **Hospital (n (%))** | | | |
| *Norfolk and Norwich* | - | 101 (100.0) | |
| *St Mary's* | 65 (48.9) | - | |
| *Queen Charlotte's* | 54 (40.6) | - | |
| *Addenbrookes* | 14 (10.5) | - | |
| **Total diet composition (n (%))** | | | |
| *Breast milk* | 36 (27.1) | 70 (69.3) | <0.001 |
| *Breast milk + Donor breast milk* | 68 (51.1) | 7 (6.9) | |
| *Breast milk + Formula* | 13 (9.8) | 14 (13.9) | |
| *Breast milk + Donor breast milk + Formula* | 13 (9.8) | 5 (5.0) | |
| *Donor milk* | 0 (0.0) | 1 (1.0) | |
| *Formula* | 3 (2.3) | 4 (4.0) | |

n = number of infants.

SD = standard deviation.

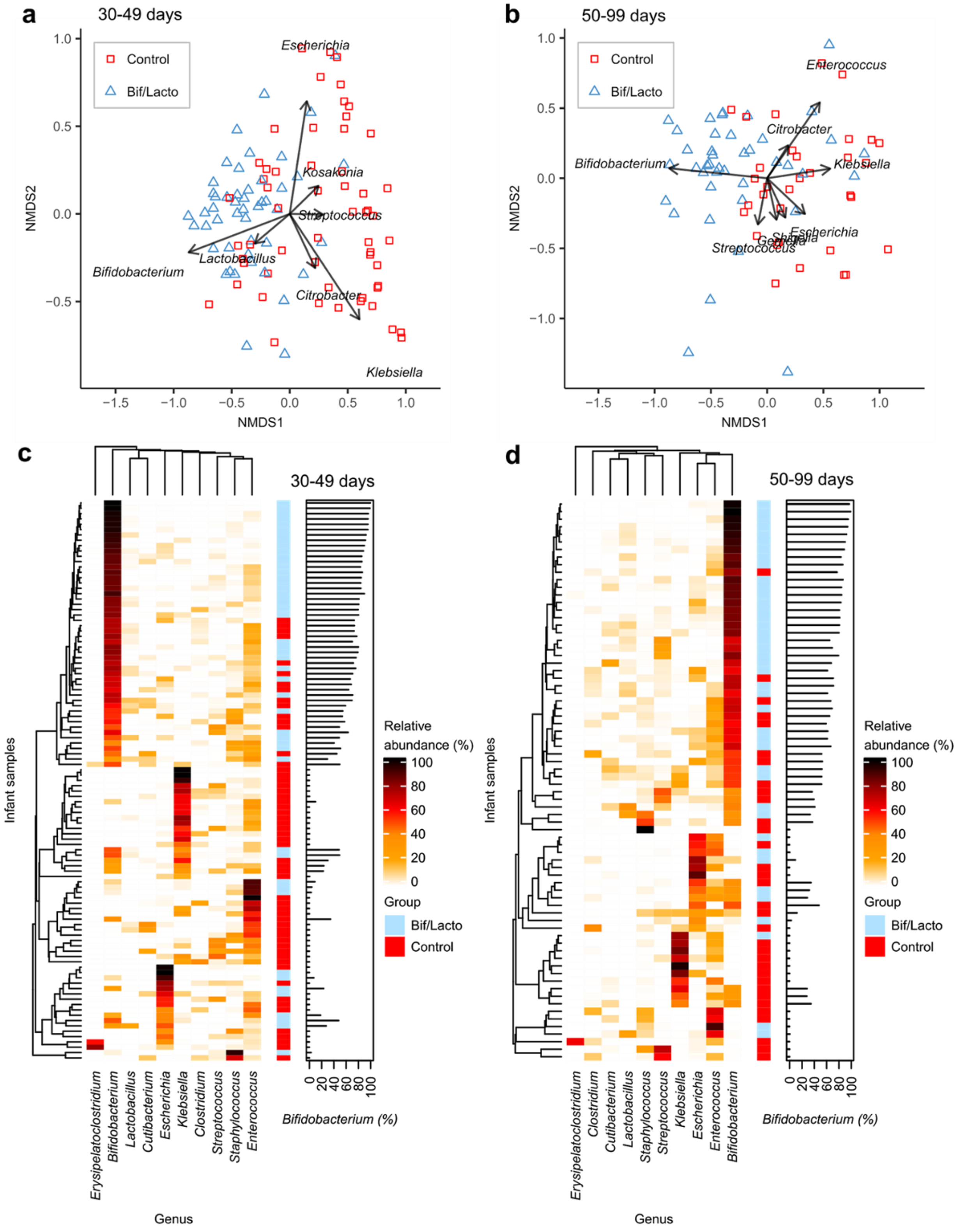


**Supplementary Figure 1: Clustering of samples and genus composition.**
**a**, Infant fecal microbiota similarity at 30-49 days and **b**, at 50-99 days shown using NMDS (non-metric multidimensional scaling) analysis clustered with a Bray-Curtis dissimilarity. Arrows indicate bacterial genera driving the separation of points on the NMDS plots. **c**, Heatmaps showing the ten genera with highest proportional abundance at 30-49 days and **d**, at 50-99 days of age. Heatmap rows were clustered using Bray-Curtis dissimilarity. Side bar plots show the proportional abundance of *Bifidobacterium* in each sample.

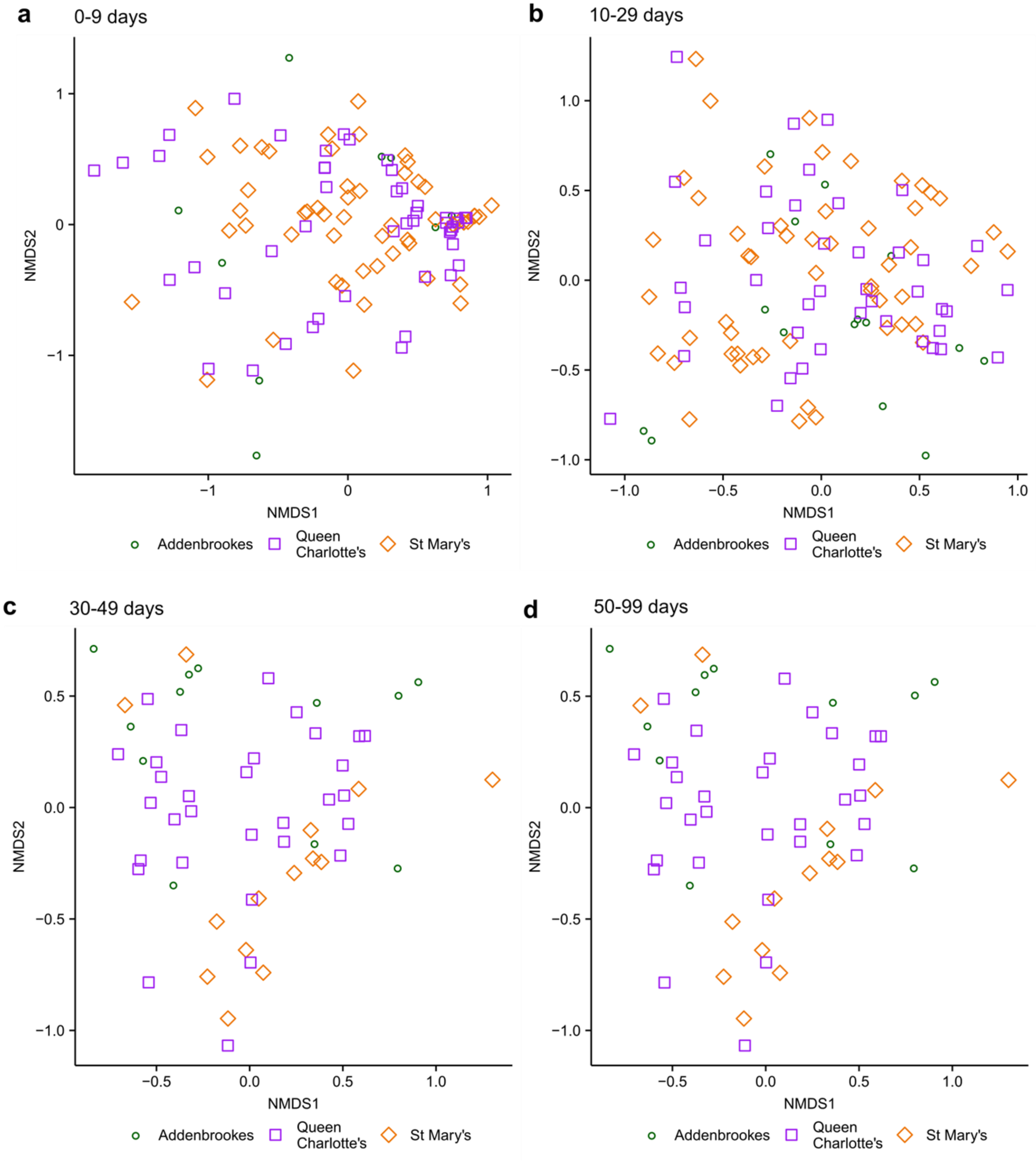


**Supplementary Figure 2: Non-metric multidimensional scaling (NMDS) comparing microbiota composition between Control hospital NICUs.**
**a**, Infant fecal microbiota composition compared by NICU at 0-9 days (Stress: 0.16; PERMANOVA: P = 0.42). **b**, Infant fecal microbiota composition compared by NICU at 10-29 days (Stress: 0.19; PERMANOVA: P = 0.33). **c**, Infant fecal microbiota composition compared by NICU at 30-49 days (Stress: 0.17; PERMANOVA: P = 0.13). **d**, Infant fecal microbiota composition compared by NICU at 50-99 days (Stress: 0.17; PERMANOVA: P = 0.029, Post-hoc analysis P > 0.05). NMDS analysis clustered using Bray-Curtis dissimilarity.

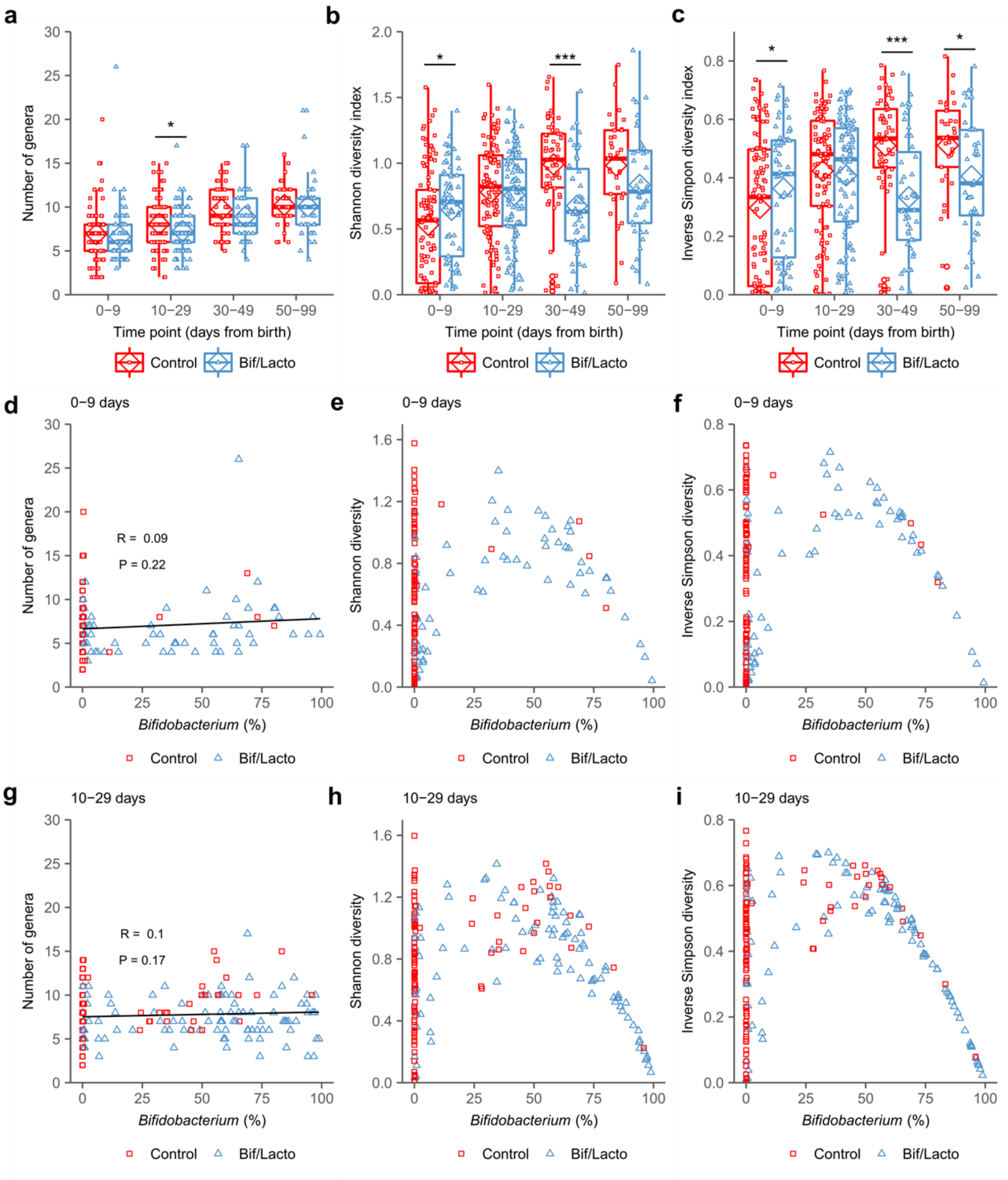


**Supplementary Figure 3: Microbiota diversity.**
**a**, Number of genera detected in each infant sample. **b**, Shannon diversity index of each infant sample. **c**, Inverse Simpson diversity index of each infant sample. **d**, Number of genera against *Bifidobacterium* abundance at 0-9 days. **e**, Shannon diversity against *Bifidobacterium* abundance at 0-9 days. **f**, Inverse Simpson diversity against *Bifidobacterium* abundance at 0-9 days. **g**, Number of genera against *Bifidobacterium* abundance at 10-29 days. **h**, Shannon diversity against *Bifidobacterium* abundance at 10-29 days. **i**, Inverse Simpson diversity *Bifidobacterium* abundance at 10-29 days. Individual points show each infant sample, the diamond indicates the group mean, box plots show median and interquartile ranges.

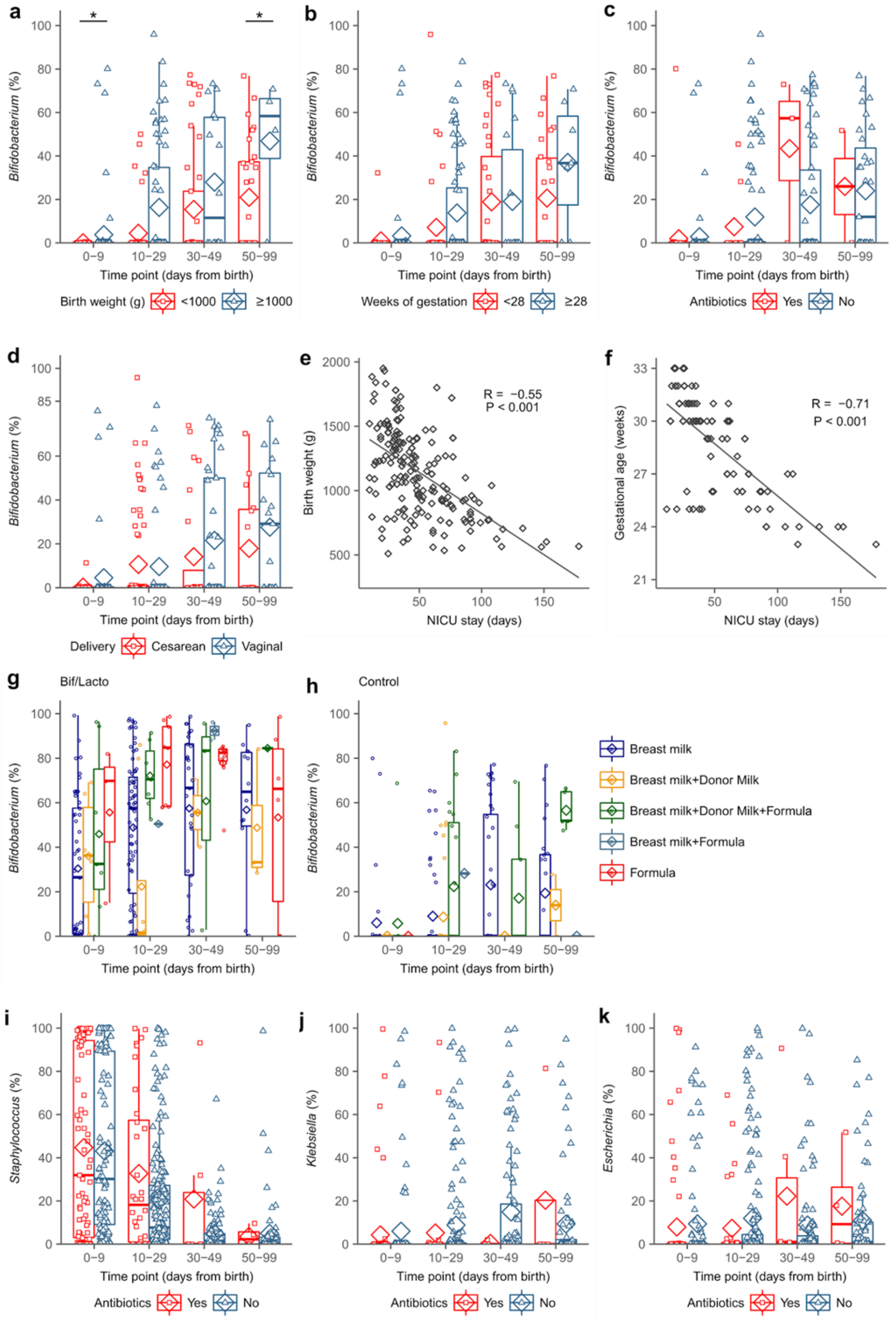


**Supplementary Figure 4: Effects of birthweight, antibiotics, delivery mode and diet on *Bifidobacterium*.**

**a**, Control group *Bifidobacterium* abundance between very low birth weight (<1000 g) and low birth weight (≥1000 g) infants. **b**, Control group *Bifidobacterium* abundance infants with very low gestational age (<28 weeks) and low gestational age (≥28 weeks). **c**, Control group *Bifidobacterium* abundance in infants receiving antibiotics at the time of sample collection. **d**, Control group *Bifidobacterium* abundance in infants delivered by caesarean and vaginal birth. **e**, Birth weight correlated with length of stay in NICU in all infants. **f**, Gestational age correlated with length of stay in NICU in all infants. **g,** Bif/Lacto group *Bifidobacterium* abundance by diet group. **h,** Control group *Bifidobacterium* abundance by diet group. **i**, *Staphylococcus* abundance in infants receiving antibiotics at the time of sample collection. **j**, *Klebsiella* abundance in infants receiving antibiotics at the time of sample collection. **k**, *Escherichia* abundance in infants receiving antibiotics at the time of sample collection. Asterisks represent *p* values: *P < 0.05.

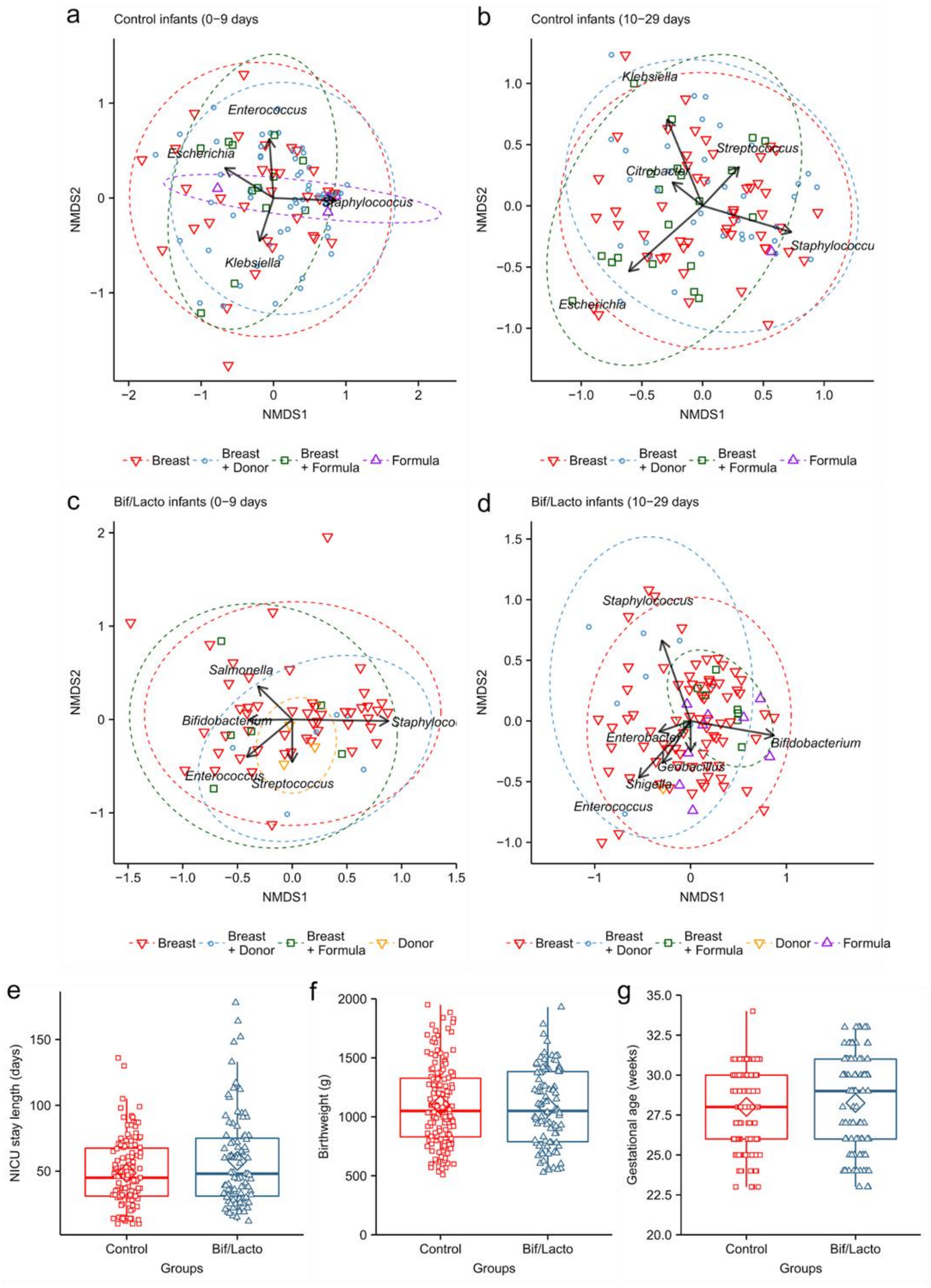


**Supplementary Figure 5: Effects of birthweight, antibiotics, delivery mode and diet on *Bifidobacterium*.**
**a**, Control infant fecal microbiota composition compared by diet at 0-9 days. **b**, Control infant fecal microbiota composition compared by diet at 10-29 days. **c**, Bif/Lacto infant fecal microbiota composition compared by diet at 0-9 days. **d**, Bif/Lacto infant fecal microbiota composition compared by NICU at 10-29 days. NMDS analysis clustered using Bray-Curtis dissimilarity. Ellipses show 95% confidence interval for each group. **e**, Length of infant stay in NICU. **f**, Infant birth weight. **g**, Infant gestational age at birth. Box plots show mean (diamond) and median (solid line) with point showing individual infants.

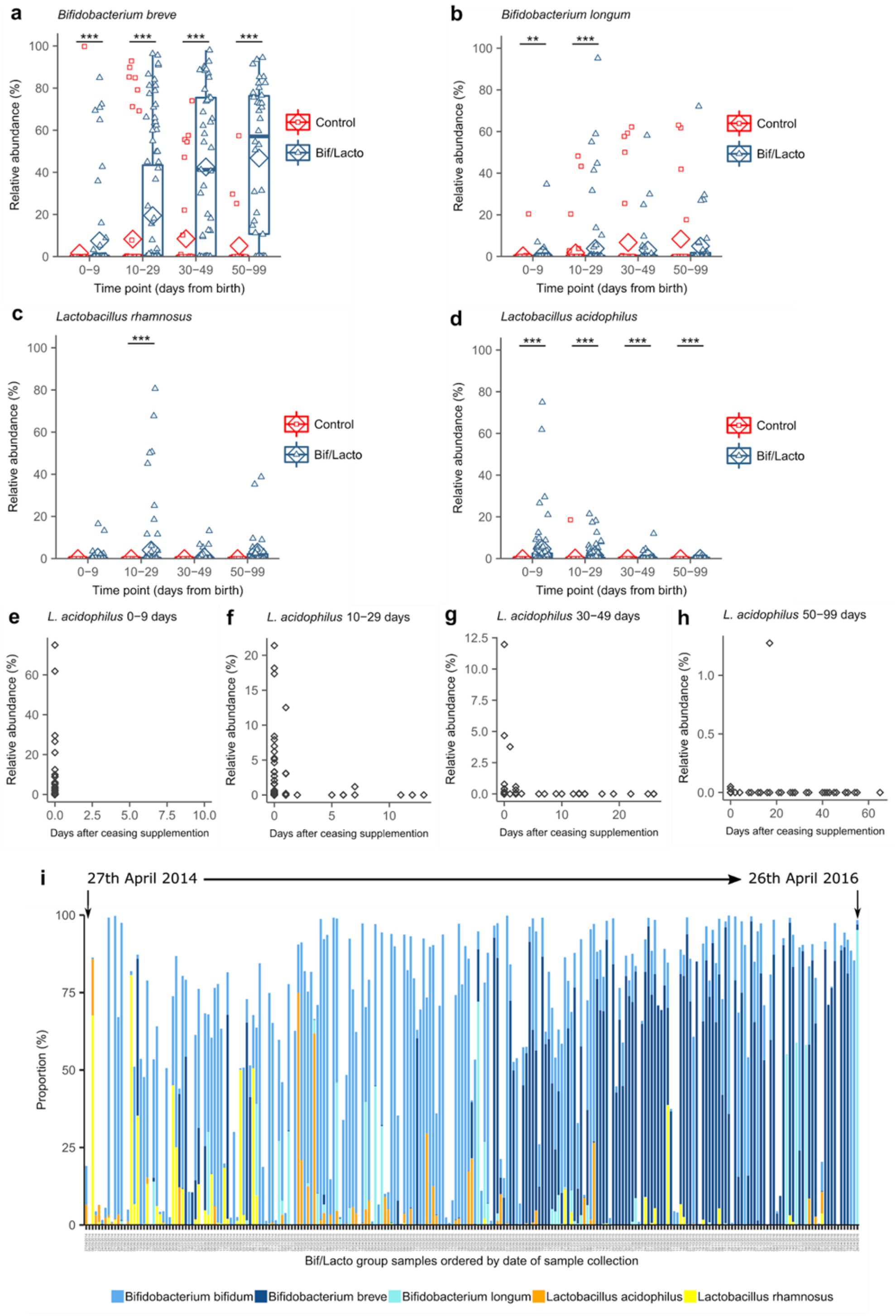


**Supplementary Figure 6: *Bifidobacterium* species level abundance and over time.**
**a,** *Bifidobacterium breve* proportional abundance. **b,** *Bifidobacterium longum* proportional abundance. **c**, *Lactobacillus rhamnosus* proportional abundance. **d**, *Lactobacillus acidophilus* proportional abundance. **e-h,** Correlation between *Lactobacillus acidophilus* abundance and days after ceasing receiving supplementation. **i**, *Bifidobacterium* and *Lactobacillus* species abundance in Bif/Lacto infant samples arranged in chronological order by date of sample collection. Asterisks represent *p* values: *P < 0.05, **P < 0.01, ***P < 0.001.

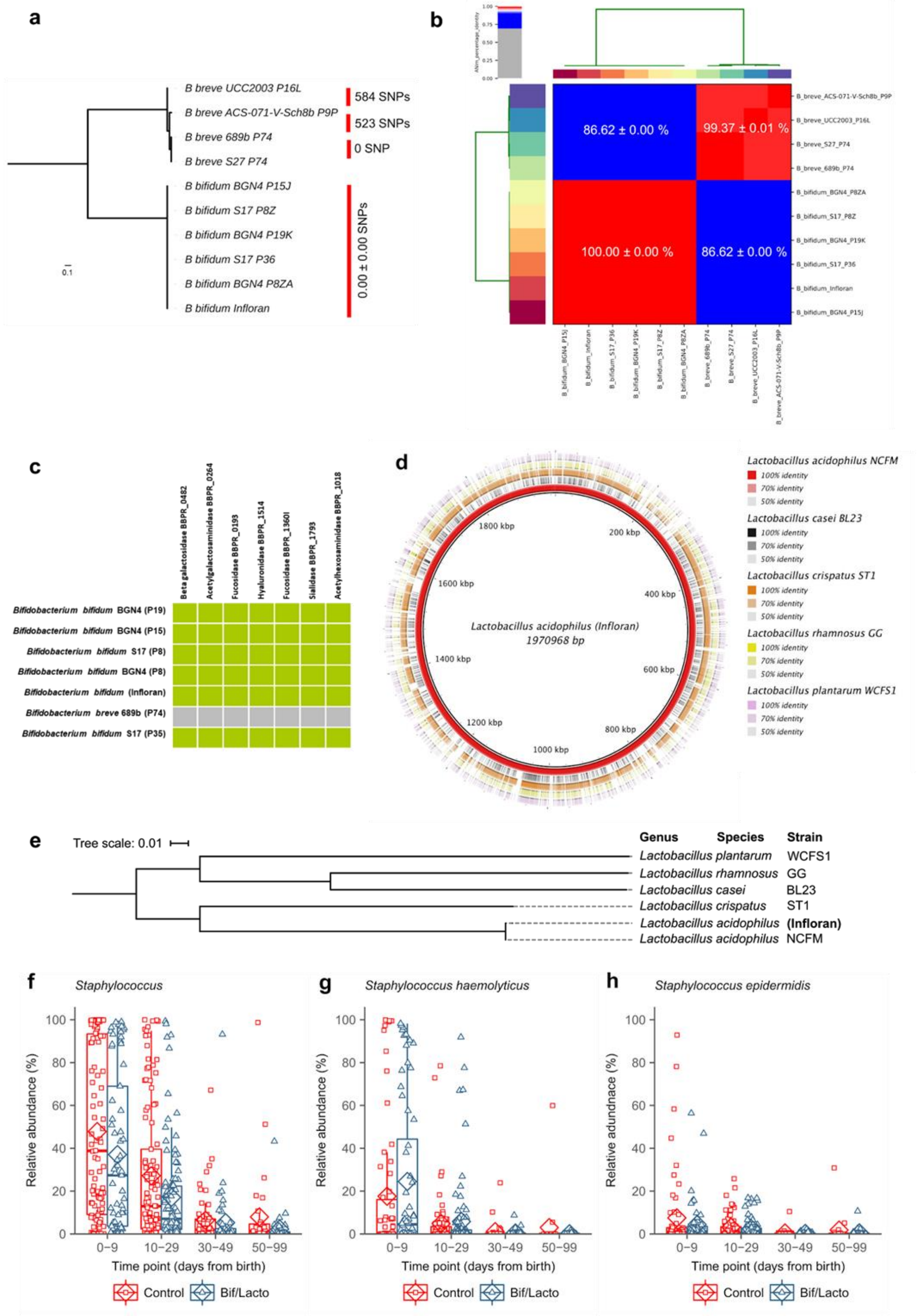


**Supplementary Figure 7: *Lactobacillus* genome comparison and *Staphylococcus* species.**
**a**, Mid-point rooted maximum-likelihood tree based on 6,202 SNPs in 87 core genes from 10 *Bifidobacterium* genomes. Data: mean ± S.D. **b**, Average Nucleotide Identity pairwise comparison between 10 *Bifidobacterium* strains. Data: mean ± S.D. **c**, Heat map representing *B. bifidum* genes involved in mucin degradation. **d**, Circular genome diagrams from *Lactobacillus acidophilus* present in the oral supplementation and a subset of five *Lactobacillus* species from NCBI database. Similarity was calculated using BLAST. **e**, Core genome tree comparison from *Lactobacillus* present in the oral supplementation and five other *Lactobacillus* species from NCBI database which were found most abundant in the 16S rRNA gene data. Roary core gene alignment output was used to create a maximum likelihood (ML) phylogenetic tree **f**, *Staphylococcus* genus relative abundance. **g**, *Staphylococcus haemolyticus* relative abundance. **h**, *Staphylococcus epidermis* relative abundance.

**a**

| Timepoint | n | Q2Y | p |
|---|---|---|---|
| 1 | 38 | 0.562(1) | 0.01 |
| 2 | 44 | 0.652(0) | 0.01 |
| 3 | 44 | 0.520(1) | 0.01 |
| 4 | 24 | 0.465(0) | 0.01 |
| Combined | 157 | 0.587(1) | 0.01 |

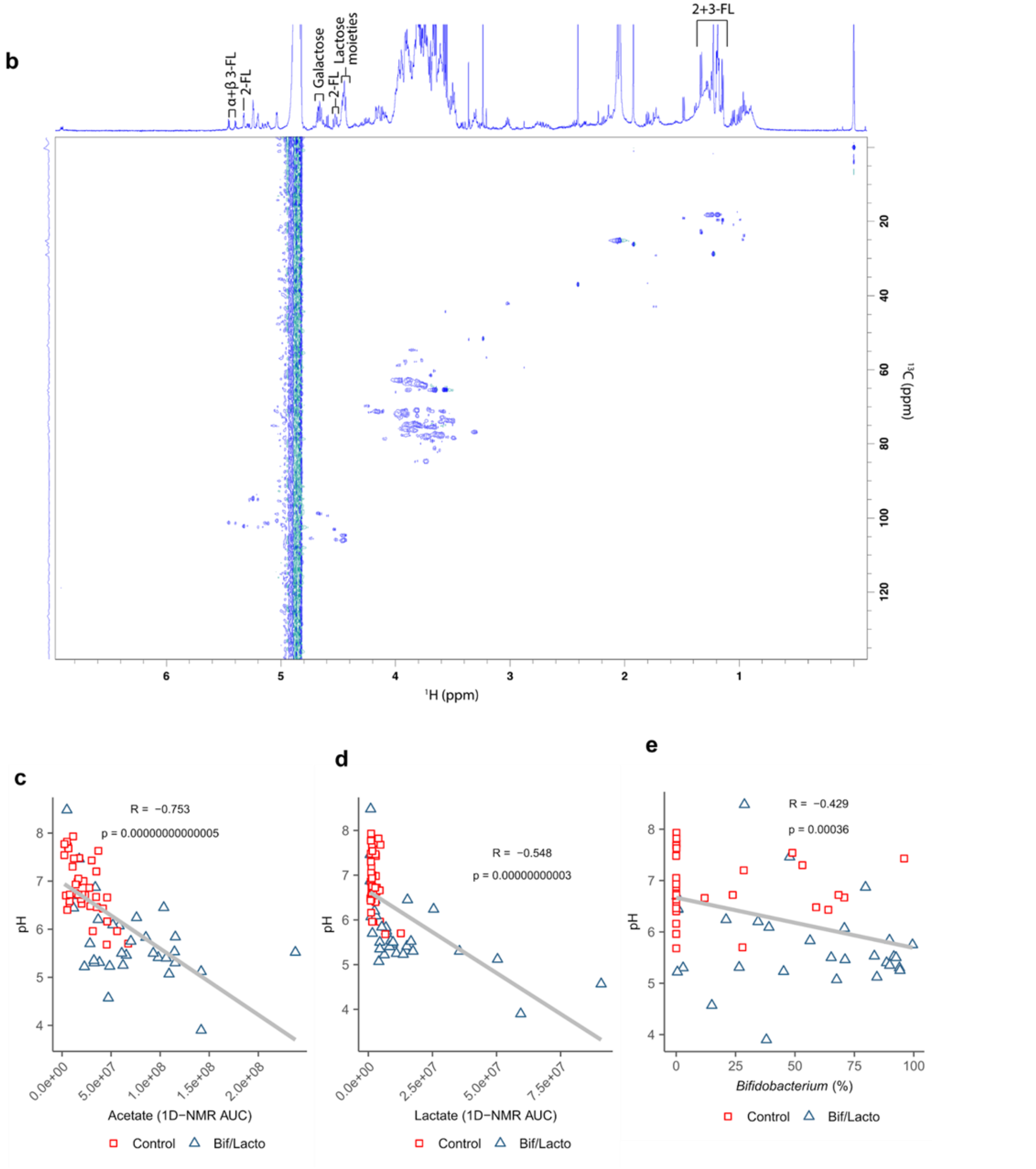


**Supplementary Figure 8: OPLS-DA model comparing the fecal ¹H NMR spectra of the Bif/Lacto Group and Control Group, and 2D-NMR analysis**
**a**, Predictive performance (Q2Y) and *p* values of the Orthogonal Projections to Latent Structures Discriminant Analysis (OPLS-DA) models comparing the Bif/Lacto and Control fecal profiles at individual time points and all timepoints combined. **b**, HSQC 2D-NMR spectrum from a study fecal sample. **c**, Spearman correlation between fecal acetate and fecal pH. **d**, Correlation between fecal lactate and fecal pH. **e**, Correlation between percentage of *Bifidobacterium* and fecal pH.

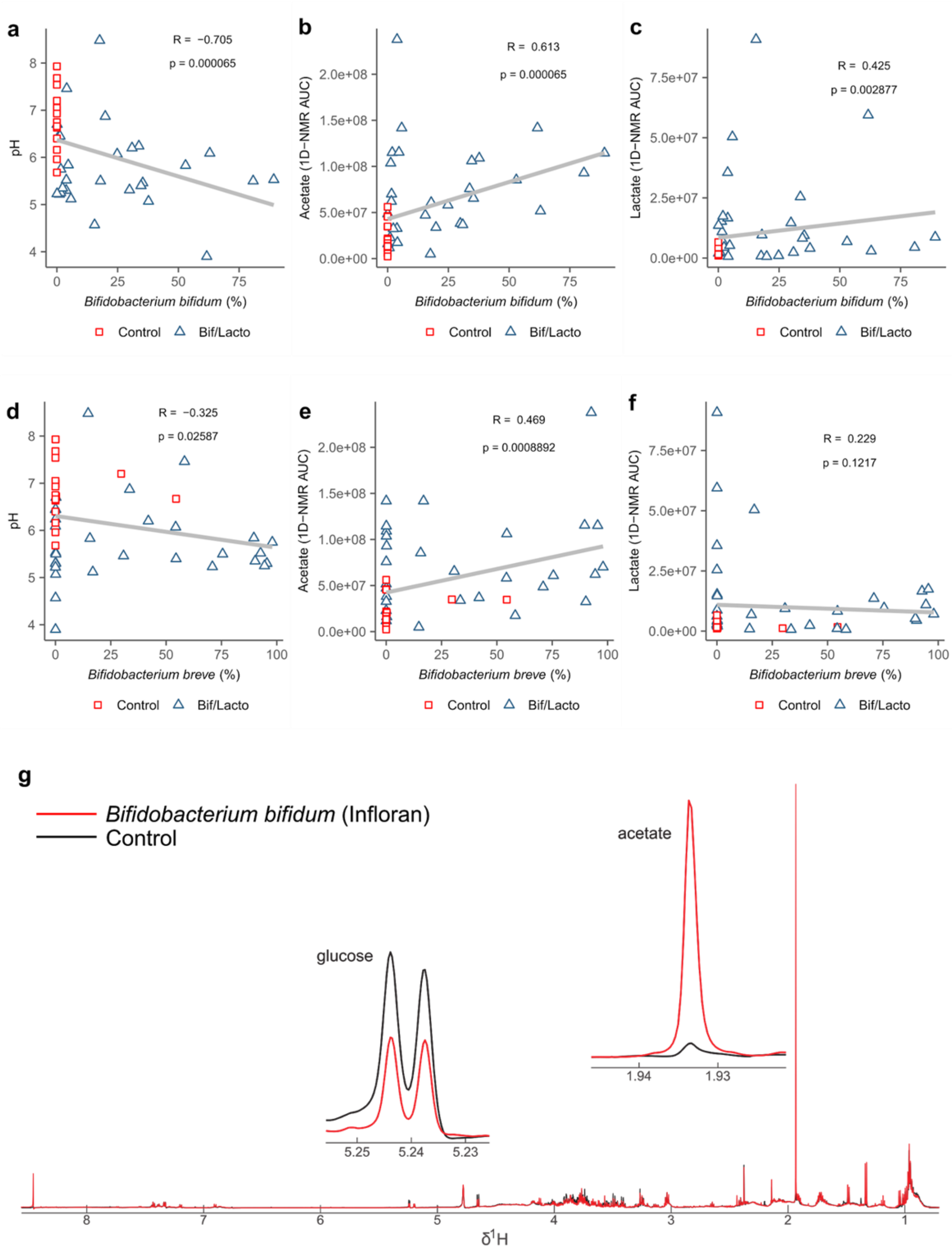


**Supplementary Figure 9: $^{1}$H-NMR AUC acetate levels found in Bif/Lacto Group and Control Group**.

**a**, Correlation between relative abundance of *Bifidobacterium bifidum* and fecal pH. **b**, Correlation between relative abundance of *B. bifidum* and fecal acetate. **c**, Correlation between relative abundance of *B. bifidum* and fecal lactate. **d**, Correlation between relative abundance of *Bifidobacterium breve* and fecal pH. **e**, Correlation between relative abundance of *B. breve* and fecal acetate. **f**, Correlation between relative abundance of *B. breve* and fecal lactate. **g**, $^{1}$H NMR spectra of the culture media from *Bifidobacterium bifidum* Infloran (red) and negative control (black).
